# High-Throughput Site-Specific N-Glycosylation Profiling of Human Fibrinogen in Atrial Fibrillation

**DOI:** 10.1101/2025.01.29.635427

**Authors:** Dinko Šoić, Domagoj Kifer, Janko Szavits-Nossan, Aleksandar Blivajs, Lovorka Đerek, Diana Rudan, Olga Gornik, Ivan Gudelj, Toma Keser

**Author notes:** Correspondence: Toma Keser, Ivan Gudelj.

## Abstract

Fibrinogen is a major plasma glycoprotein involved in blood coagulation and inflammatory responses. Alterations in its glycosylation have been implicated in various pathological conditions, yet its site-specific N-glycosylation profile remains largely unexplored in a clinical context. Here, we present a high-throughput LC-MS workflow for site-specific analysis of fibrinogen N-glycosylation using a cost-effective ethanol precipitation enrichment method. The method demonstrated good intra- and inter-plate repeatability (CV: 5% and 12%, respectively) and was validated through the first assessment of intraindividual temporal stability in healthy individuals, revealing consistent glycosylation patterns within individuals. Application to 181 atrial fibrillation (AF) patients and 52 healthy controls identified three gamma chain glycoforms significantly associated with AF. Most notably, increased levels of the asialylated N4H5, known to enhance fibrin bundle thickness and promote clot formation, suggest a potential mechanism linking glycosylation changes to the prothrombotic state in AF. Furthermore, fibrinogen sialylation showed strong associations with cardiovascular risk factors, including triglycerides, BMI, and glucose levels. Longitudinal analysis of 108 AF patients six months post-catheter ablation showed stability in the AF-associated glycan profile. Our findings establish fibrinogen glycosylation as a potential biomarker for cardiovascular conditions and demonstrate the utility of site-specific glycosylation analysis for clinical applications.

## Introduction

Fibrinogen is a 340 kDa plasma glycoprotein exhibiting multiple homeostasis and inflammatory functions beyond its classical role in blood coagulation (1,2). It is an acute-phase protein synthesized in hepatocytes and one of the most abundant glycoproteins in blood plasma at concentrations of 1.5-4.5 g/L (3). Its hexameric structure comprises two sets of three polypeptide chains (Aα, Bβ, γ) arranged in a symmetrical dimeric configuration (3). This complexity hosts distinct binding domains and allows diverse interaction capabilities, supporting its multifunctional nature (1). The strategic position of fibrinogen’s N-glycans on its Bβ and γ chains provides an opportunity to monitor disease-specific modifications that could influence both its coagulation and inflammatory functions (4). While glycosylation changes of other abundant plasma proteins, notably immunoglobulin G (5), have emerged as powerful indicators of various pathological states , the diagnostic potential of fibrinogen glycosylation remains largely unexplored.

Fibrinogen’s complex structure and extensive post-translational modifications underlie its diverse biological roles, facilitating multiple critical interactions (4). During coagulation, thrombin-mediated cleavage of fibrinogen initiates polymerization of fibrin monomers, leading to the formation of a cross-linked network which provides the structural foundation for thrombus formation (6). Furthermore, interaction with platelet integrin αIIbβ3 enables platelet aggregation (2,7). Beyond hemostasis, fibrinogen supports wound healing through epithelial regeneration, and interacts with endothelial cells via VE-cadherin to influence vascular function and angiogenesis (2,8). Immune responses are modulated by engaging with leukocyte integrin αMβ2 (Mac-1), triggering NF-κB pathway activation and subsequent inflammatory cytokine production (1,2,9). Elevated fibrinogen plasma levels during acute-phase responses to tissue injury or inflammation also significantly contribute to innate immunity (2,10,11).

Experimental evidence from genetic and pharmacological studies demonstrates fibrinogen’s causative role in various inflammatory conditions (2). Fibrinogen-deficient mice exhibit reduced inflammation in models of arthritis (12), colitis (13), and muscular dystrophy (14), among others (2). Similarly, targeting fibrinogen-receptor interactions through specific peptide inhibitors or genetic modification in mice significantly ameliorates inflammatory pathology (9). In cardiovascular disease, elevated fibrinogen levels enhance blood viscosity, endothelial activation, and leukocyte recruitment, contributing to atherosclerotic progression (15–18).

Among the various post-translational modifications fine-tuning fibrinogen’s interactions, glycosylation emerges as a particularly significant modulator of its activity (4). Glycosylation is an enzymatically regulated process where complex carbohydrate structures, known as glycans, are attached to specific amino acid residues on proteins (19). Fibrinogen undergoes both O- and N-glycosylation. Of its four potential N-glycosylation sites, only Asn394 of the Bβ chain and Asn78 of the γ chain carry N-glycans, while the two potential sites on the Aα chain remain unoccupied (20,21). The N-glycan profile is dominated by biantennary structures, with digalactosylated monosialylated (A2G2S1) and disialylated (A2G2S2) glycoforms comprising approximately 90% of the total glycoforms, whereas structures containing bisecting N-acetylglucosamine or core fucosylation appear in minor amounts (20,22). Additionally, fibrinogen undergoes extensive O-glycosylation at multiple sites on the Aα chain and a single site on the Bβ chain, though these modifications occur at substoichiometric levels, with non-glycosylated forms predominating by several orders of magnitude (22).

Multiple studies have highlighted the significance of fibrinogen N-glycosylation in determining the structural and functional properties of the protein. For instance, the sialic acid content modulates fibrin polymerization through electrostatic effects - increased sialylation inhibits fibrin polymerization by enhancing repulsion between fibrin monomers, directly impacting clot formation dynamics. Desialylated fibrinogen thus produces thicker fibrin fibers and exhibits shorter thrombin clotting times, potentially increasing thrombotic risk (23). The overall level of sialylation may also affect fibrinogen’s solubility, showcasing its impact on blood clotting (24,25). Similarly, non-enzymatic glycation of fibrinogen, particularly prevalent in diabetic patients, leads to fibrin structures more resistant to plasmin-mediated degradation *in vitro*, potentially contributing to increased cardiovascular complications (26).

These structurally significant glycan modifications have emerged as potential disease indicators, particularly in conditions affecting cardiovascular health. In end-stage renal disease (ESRD), distinct aberrations in fibrinogen glycosylation have been observed, characterized by increased levels of multi-antennary N-glycans and altered fucosylation patterns (27). Age-related changes in fibrinogen glycosylation have also been documented, with older individuals exhibiting distinct glycan patterns that may influence their susceptibility to thrombotic events (28). Furthermore, cirrhosis significantly alters fibrinogen’s glycosylation patterns, which may contribute to coagulation abnormalities observed in advanced liver disease (29).

While these disease associations are promising, their full characterization has been constrained by analytical limitations. Traditional lectin-based approaches, though valuable, provide only semi-quantitative assessment of specific glycan motifs and lack the ability to deliver comprehensive structural characterization. The inherent selectivity limitations, limited to specific glycan motifs recognized by lectins and often introducing measurement biases, combined with potential cross-reactivity and the inability to distinguish glycoforms at specific attachment sites, highlight the need for more sophisticated analytical strategies (30,31). Mass spectrometry (MS)-based glycoproteomics offers a powerful alternative, enabling site-specific glycosylation analysis with detailed glycan structural information. In recent years, numerous LC-MS glycoproteomics workflows for human plasma glycoproteins have been developed, demonstrating their broad applicability for biomarker discovery across various diseases (32–35). Although previous MS studies have characterized fibrinogen N-glycosylation using commercially available purified protein (21,22), a robust methodology for analyzing fibrinogen glycosylation directly from clinical plasma samples - essential for biomarker discovery - has remained elusive.

Thus, we have developed a novel, cost-effective, and high-throughput LC-MS workflow for site-specific analysis of fibrinogen N-glycosylation at the intact glycopeptide level. Our method employs a straightforward ethanol precipitation approach for fibrinogen enrichment from plasma, eliminating the need for expensive antibody-based purification while maintaining sufficient specificity for reliable glycopeptide analysis. The method’s robustness and clinical applicability were validated through the first systematic assessment of intraindividual temporal stability of fibrinogen N-glycosylation in healthy individuals. Importantly, our workflow achieves the throughput and reproducibility necessary for large-scale clinical studies while providing detailed structural information about individual glycoforms at each site.

To demonstrate the clinical utility of our workflow, we conducted a comprehensive study of atrial fibrillation (AF), analyzing samples from 181 AF patients and 52 healthy controls. This application is particularly relevant given that AF, the most common cardiac arrhythmia, has complex relationships with coagulation and inflammation - processes in which fibrinogen plays central roles (36). Previous studies have established associations between elevated levels of coagulation factors and both the prevalence and incidence of AF, suggesting perturbations in the coagulation system may contribute to AF pathogenesis (37,38). Although altered fibrinogen glycosylation patterns have been linked to cardiovascular risk in lectin studies, no previous investigation has examined fibrinogen glycosylation changes specifically in AF. Our study not only characterized site-specific N-glycan alterations in AF patients but also examined their relationships with anthropometrical and biochemical parameters. Additionally, we tracked glycosylation changes during a six-month follow-up period after catheter ablation, providing the first longitudinal assessment of fibrinogen glycosylation in relation to AF recurrence.

## Experimental procedures

### Experimental design and statistical rationale

The manuscript comprises three sets of experiments: method development, intra-individual temporal stability analysis, and a pilot study on individuals with atrial fibrillation (AF).

In the method development phase, the analytical method was developed and optimized using a blood plasma standard derived from pooled plasma samples from the AF population. The method’s repeatability was evaluated by calculating the coefficient of variation (CV) from technical replicates of the pooled plasma standard. Intra-plate repeatability was assessed using 8 replicates, randomized across a single 96-well plate, while inter-plate repeatability was determined using 16 replicates randomized across four different 96-well plates.

All glycoforms identified and annotated in the pooled plasma standard were included in the quantification and subsequent data analysis for both the intra-individual temporal stability and AF pilot studies. Extracted signals were summed and normalized to the total integrated area for each glycosylation site. Normalization was applied to eliminate variations in signal intensity between samples, thereby enabling their comparison.

The intra-individual temporal stability study aimed to assess the stability of fibrinogen glycosylation patterns over time in healthy individuals. Samples were collected from 14 age-matched healthy male participants at three time points (0, 6, and 10 weeks). All samples were randomized across a 96-well plate prior to analysis. For each participant, the intra-individual CV was calculated from their longitudinal samples, while the inter-individual CV was calculated from all participants’ samples at each time point.

The AF pilot study included a total of 233 subjects: 181 individuals with AF who underwent catheter ablation and 52 cardiovascularly healthy controls. Among the 181 AF patients, 108 were sampled again six months after ablation. All samples were randomized across four 96-well plates before analysis.

Data analysis and visualization were performed using the R programming language (version 4.3.3) and Microsoft Excel 2016 (Microsoft Corp).

Batch effects were corrected using an empirical Bayesian framework (R package sva) (39). The batch-adjusted data were used for all subsequent analyses.

The effects of sex and age on the levels of each glycopeptide were estimated using a mixed-effects model. Glycoproteomic abundance was set as the dependent variable, with sex and age included as fixed factors. Subject identifiers, as well as group and timepoint (nested within the group), were modeled as random intercepts (R package lme4) (40). Before analysis, glycopeptide levels were transformed using an inversion rank transformation to approximate a standard normal distribution (rankit transformation) (41).

To compare groups and assess changes before and after catheter ablation, a mixed-effects model was applied, with rankit-transformed glycopeptide levels as the dependent variable. Group and timepoint (nested within the group) were included as fixed factors, while sex and age were added as covariates. Subject identifier was modeled as a random intercept. Post-hoc tests (R package emmeans) were conducted to estimate differences in glycopeptide levels: (i) before and after ablation and (ii) between control and case groups (pre-ablation) (42).

To evaluate differences between subjects with and without recurrence after ablation, a mixed-effects model was utilized. Rankit-transformed glycopeptide levels were the dependent variable, with recurrence status (Yes/No) crossed with timepoint as a fixed factor. Subject identifier was included as a random intercept. Post-hoc tests were performed to compare glycopeptide levels between recurrence groups at each timepoint and to assess the before-to-after difference (difference of differences).

Relationships between glycopeptide levels and blood parameters, biochemical data, or body mass index (BMI) were analyzed using mixed-effects or linear models, depending on data availability. Each model included rankit-transformed glycopeptide levels as the dependent variable and the rankit-transformed variable of interest as a predictor. Sex and age were included as covariates, with group, timepoint (nested within the group), and subject identifier added as random intercepts, if applicable.

For all analyses, p-values were adjusted for multiple testing using the false discovery rate (Benjamini-Hochberg method, modified by Li and Ji) (43,44). The level of statistical significance was set at α = 0.05.

### Population for the intra-individual temporal stability study

Fourteen male physical education students (age 19 ± 0.7 years) participated in the study. All participants were screened for cardiovascular diseases, muscle injuries or ongoing medical treatment before their inclusion into the experimental protocol. Participants were instructed to refrain from alcohol and cigarette consumption as well as antioxidant supplementation throughout the study. The blood samples were taken from each participant at three time points: 0, 6 and 10 weeks. The blood was collected in vacuum tubes containing EDTA with 20-G straight needle venipuncture from the antecubital vein. The EDTA tubes were immediately centrifuged (at 1370g for 10 minutes) to separate erythrocytes from plasma. Subsequently, plasma supernatant was aspirated into a series of 1 mL aliquots and stored at - 80 °C until analysis.

### Population for the atrial fibrillation study

The study population was described in detail previously (45,46). Briefly, this study included a total of 233 individuals: 181 patients with paroxysmal or persistent AF indicated for pulmonary vein isolation via radiofrequency catheter ablation, and 52 healthy controls. Plasma samples of AF patients were collected at Magdalena Clinic, Krapinske Toplice, while samples from healthy controls were collected at Dubrava Clinical Hospital, Zagreb. Venous blood samples from all participants were collected in vacuum tubes containing tri-potassium ethylenediaminetetraacetic acid (K3EDTA). The samples were allowed to rest for an hour, and then centrifuged at 1620 g for 10 minutes. Plasma aliquots were transferred to 2 mL tubes, centrifuged again at 2700 g for 10 minutes, and immediately stored at -20°C until analysis. For AF patients, plasma samples were collected at two time points: before the procedure and at the six-month follow-up. Their demographic data are summarized in Table S1.

### Ethics Statement

Both studies are designed in accordance with the Declaration of Helsinki, supported by written informed consent from all individuals and approvals from eligible local Ethics Committees - Ethical Committee of the University of Zagreb Faculty of Pharmacy and Biochemistry, Ethical Committee of Magdalena Clinic and Ethical Committee of Dubrava Clinical Hospital.

### Enrichment of fibrinogen from plasma samples

Fibrinogen was enriched by precipitation with absolute ethanol (47). 20 μl of human plasma in each well was mixed with 3 μL of cold absolute ethanol (Merck, Darmstadt, Germany) in a 96-well PCR plate (Thermo Scientific, Rockford, IL, USA). The plate was incubated for 2 minutes at -20°C and then centrifuged for 20 minutes at 4000g at 4°C. The supernatant was discarded to an empty PCR plate using in-house 3D printed adapters in a high-throughput manner by low-speed centrifugation at 15g for 1 minute.

### Reduction, alkylation and trypsin digestion

The remaining precipitate, containing enriched fibrinogen, was resuspended by adding 45 μl of 15mM dithiothreitol (Sigma-Aldrich, St Louis, MO, USA) + 100mM ammonium bicarbonate (Acros Organics, Geel, Belgium) and the samples were incubated for 30 minutes at 60°C. After cooling to room temperature, 2 μl of 700mM iodoacetamide (Sigma-Aldrich, St Louis, MO, USA) was added and the samples were incubated in dark for 30 minutes. Subsequently, 2 μL of 0.4 μg/μL TPCK-treated trypsin (Promega, Madison, WI, USA) in 50mM acetic acid was added to each sample and they were incubated overnight at 37°C.

### Glycopeptide enrichment with HILIC-SPE

Glycopeptides were enriched using a hydrophilic interaction chromatography based solid-phase extraction (HILIC-SPE) on a 96-well polypropylene filter plate (Orochem, Naperville, IL, USA). 5 mg of Chromabond® HILIC beads (Macherey-Nagel, Düren, Germany) in 0.1% TFA in water (Sigma-Aldrich, St Louis, MO, USA) (50mg/mL suspension) was added to each well. Solvent was removed by application of vacuum using a vacuum manifold (Millipore Corporation, Billerica, MA, USA). All wells were prewashed using 2× 250 µL of 0.1% TFA in water, followed by equilibration using 2× 250 μl of 90% acetonitrile (VWR International, Radnor, PA, USA) + 10% 0.1% TFA in water. The samples were diluted with 630 µL of 0.11% TFA in acetonitrile and loaded into the wells, which were subsequently washed 2× with 250 μl of 90% acetonitrile + 10% 0.1% TFA in water. Enriched glycopeptides were eluted into a PCR plate with 200 μl of 0.1% TFA in water. The eluates were immediately dried down in a SpeedVac Vacuum Concentrator (Thermo Scientific, Rockford, IL, USA) and stored at −20°C until analysis.

### RP-LC-ESI-MS(/MS)

Separation and measurements were performed using an Acquity H-class ultra-performance liquid chromatography (UPLC) instrument (Waters, Milford, MA, USA) coupled to a Synapt G2-Si ESI-QTOF-MS mass spectrometer (Waters, Milford, MA, USA), with a Lockspray Exact Mass Ionization Source (Waters). The instrument was under the control of MassLynx v.4.1 software (Waters). Dried samples were reconstituted in 50 μL of ultrapure water and 40 μL was injected onto column. Separation of the tryptic fibrinogen glycopeptides was based on differences in their peptide backbone and was performed on a Waters bridged ethylene hybrid (BEH) C18 chromatography column, 150 × 2.1 mm, 130Å, 1.7 μm BEH particles. In the first 12 minutes solvent B (0.1% TFA in acetonitrile) was increased from 21% to 31% and during the next 1 minute from 31% to 80%, which was held for 2 minutes to wash the column. Solvent A was 0.1% TFA in water. Flow rate was 0.4 mL/min and column temperature was 30°C.

MS conditions were set as follows: positive ion mode, capillary voltage 3 kV, sampling cone voltage 40 V, source temperature 120°C, desolvation temperature 350°C, desolvation gas flow 600 l/h. Mass spectra were recorded from 500 to 2500 m/z at a frequency of 1 Hz. MS/MS experiments were performed in a data-dependent acquisition (DAD) mode. Spectra were first acquired from 500 to 2500 m/z and then two precursors with the highest intensities were selected for CID fragmentation (m/z 200 to 2000 was recorded). A collision energy ramp was used for the fragmentation (Low Mass 500 Da – CE Ramp Start 15 V, CE Ramp End 30 V; High Mass 2500 Da – CE Ramp Start 60 V, CE Ramp End 80 V). For low intensity peaks MS/MS experiments were performed in Tof MRM mode (m/z 100 to 2500 was recorded) with the same CE Ramp as in DDA.

### Proteomic Data Analysis

To assess the effectiveness of our fibrinogen enrichment protocol and identify coenriched glycoproteins in the ethanol precipitate, we conducted a proteomic analysis using open-source SearchGUI software (version 4.3.11) (48), with Sage as a chosen proteomics database search engine (49). The human reference proteome (release version 2024_10_10, ProteomeID: UP000005640_9606) with 20,412 reviewed protein sequences was obtained from Uniprot. Precursor mass tolerance was set to 0.1 Da, fragment mass tolerance to 0.1 Da, while all the other parameters were left at default settings. Enzyme was set to trypsin, with maximum of two misses. To analyze and visualize the results of the search, PeptideShaker was used. False discovery rate of 1% at peptide spectrum match (PSM) level was used. Oxidation (M) was set as a variable modification, while carbamidomethyl (C) was set as a fixed modification. Default MS2-based label-free quantification based on spectral counting was used, and relative abundance of each identified protein was expressed as percentage (Table S2).

### Glycoproteomic Data analysis

Fibrinogen glycopeptides spectra generated in the MS/MS experiments were manually annotated using MassLynx v.4.1 software (Waters), GlycoWorkbench (50) and GlycoMod (51), the latter two being operated under permissive free software licenses. All identified glycopeptides were validated through MS/MS analysis (refer to the Results section for detailed information).

Prior to the relative quantification of the glycoproteomic data, MSConvert tool (ProteoWizard version 3) was used to convert all Waters raw data into mzXML file format. LaCyTools (version 1.0.11.0.1 b.9), operated under free software license (52), was used for automated relative quantification of the MS data. Chromatograms were aligned based on the four most abundant glycopeptide signals. Targeted peak integration was performed on triply charged species. Signals were integrated to include at least 90% of the theoretical isotopic pattern. The quality control (QC) parameters for extracted data of the targeted peak integration were automatically calculated for each analyte of every sample: mass accuracy, deviation from the theoretical isotopic pattern and signal to noise ratio. Quality of the data was checked on the batch level and curation was performed in a software called GlycoDash (https://github.com/Center-for-Proteomics-and-Metabolomics/glycodash). Criteria were set at a level to ensure only quality extracted data to be further processed: mass accuracy (between −40 and 40=ppm), the deviation from the theoretical isotopic pattern (IPQ; below 25%), and the signal to noise ratio (above 9) of an integrated signal. Spectra were curated based on percentiles: glycosylation sites from samples with sum intensities of all passing analytes below the 5th percentile were excluded from further processing. Analytes were curated based on all data: if an analyte fulfills the QC criteria in 25% of spectra, then it passes curation and is used for further processing. The extracted signals were normalized to total integrated area per glycosylation site. Normalization serves the purpose of removing the variation in signal intensity between samples and allows for their comparison.

## Results and discussion

Development of high-throughput site-specific method for N-glycosylation analysis of fibrinogen To date, site-specific N-glycosylation of human fibrinogen has not been profiled in a cohort study. To address this, we adapted a previously established ethanol precipitation protocol by Qiu et al. (47) into a 96-well format, enabling efficient, high-throughput fibrinogen enrichment (Figure 1). This method proved cost-effective and scalable, with optimal LC-MS signals obtained by precipitating 20 μL of human plasma with 15% absolute ethanol (Figure S1). As fibrinogen is a large and complex protein with a molecular weight of approximately 340 kDa, it is among the least soluble of the major plasma proteins. Its size and structure make it particularly susceptible to ethanol precipitation compared to smaller, more compact proteins (47). Enriched fibrinogen was prepared for analysis by reducing disulfide bonds with dithiothreitol, followed by alkylation with iodoacetamide and trypsin digestion. To further purify the sample and selectively remove non-glycosylated, hydrophobic peptides, hydrophilic interaction chromatography-based solid-phase extraction (HILIC-SPE) was applied post-digestion. According to analysis of proteomics data acquired by LC-MS fibrinogen accounted for an average of 33% of all proteins in the enriched fractions (Table S2). It is worth noting that our method is not restricted to N-glycosylation analysis, as it can easily be adapted, through appropriate liquid chromatography modifications, to study other hydrophilic post-translational modifications such as O-glycosylation and phosphorylation.

**Figure 1.**
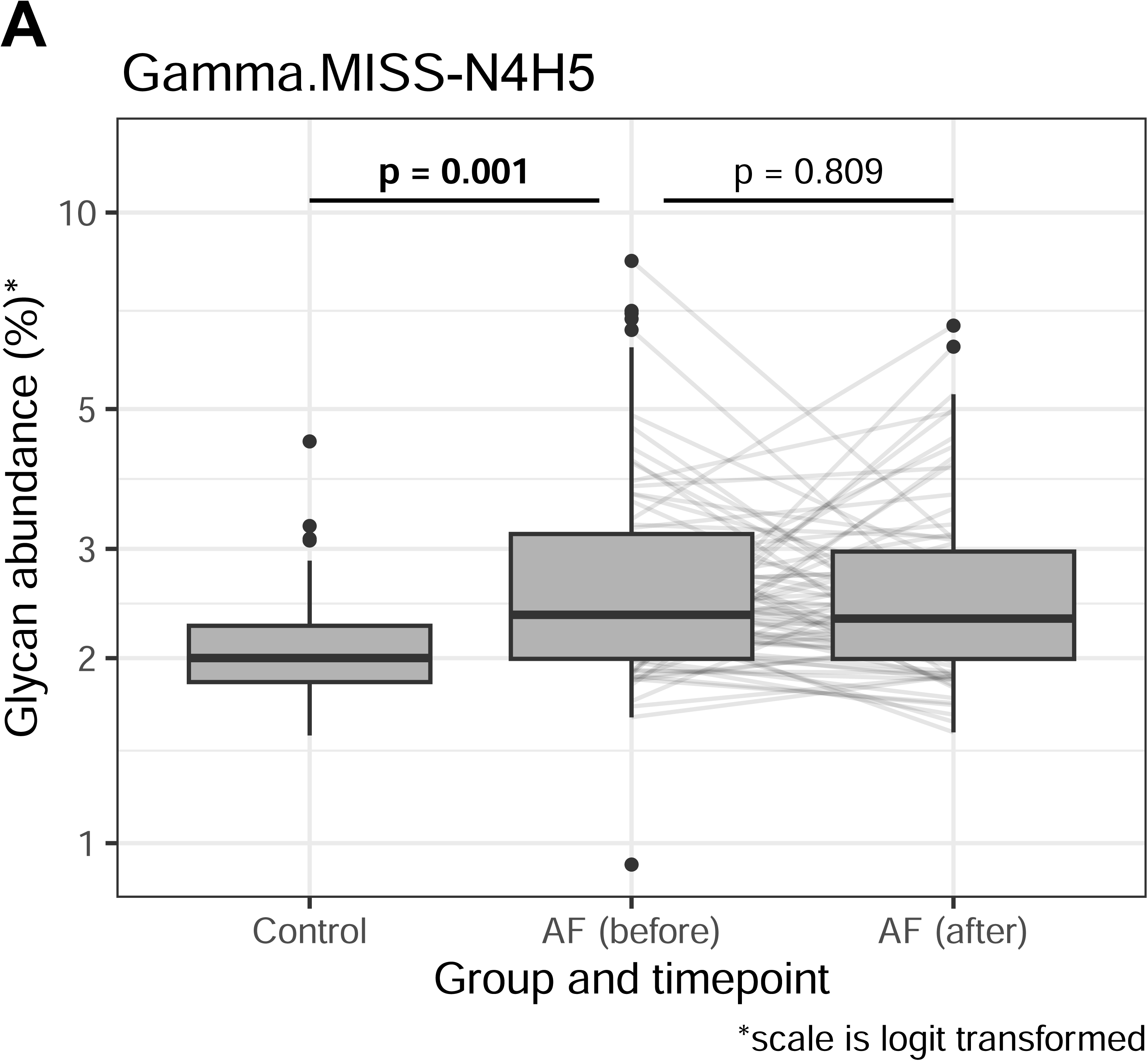
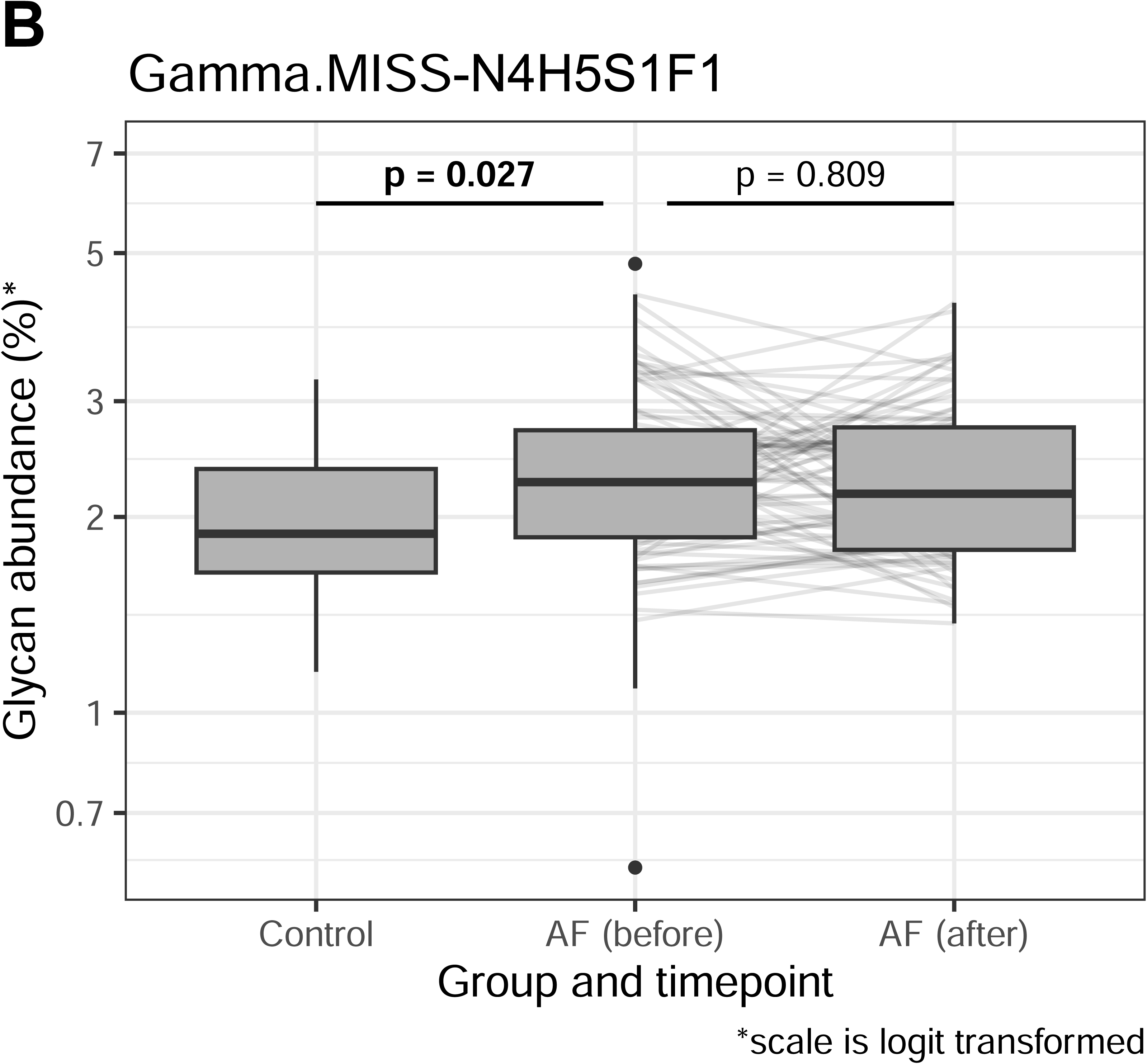
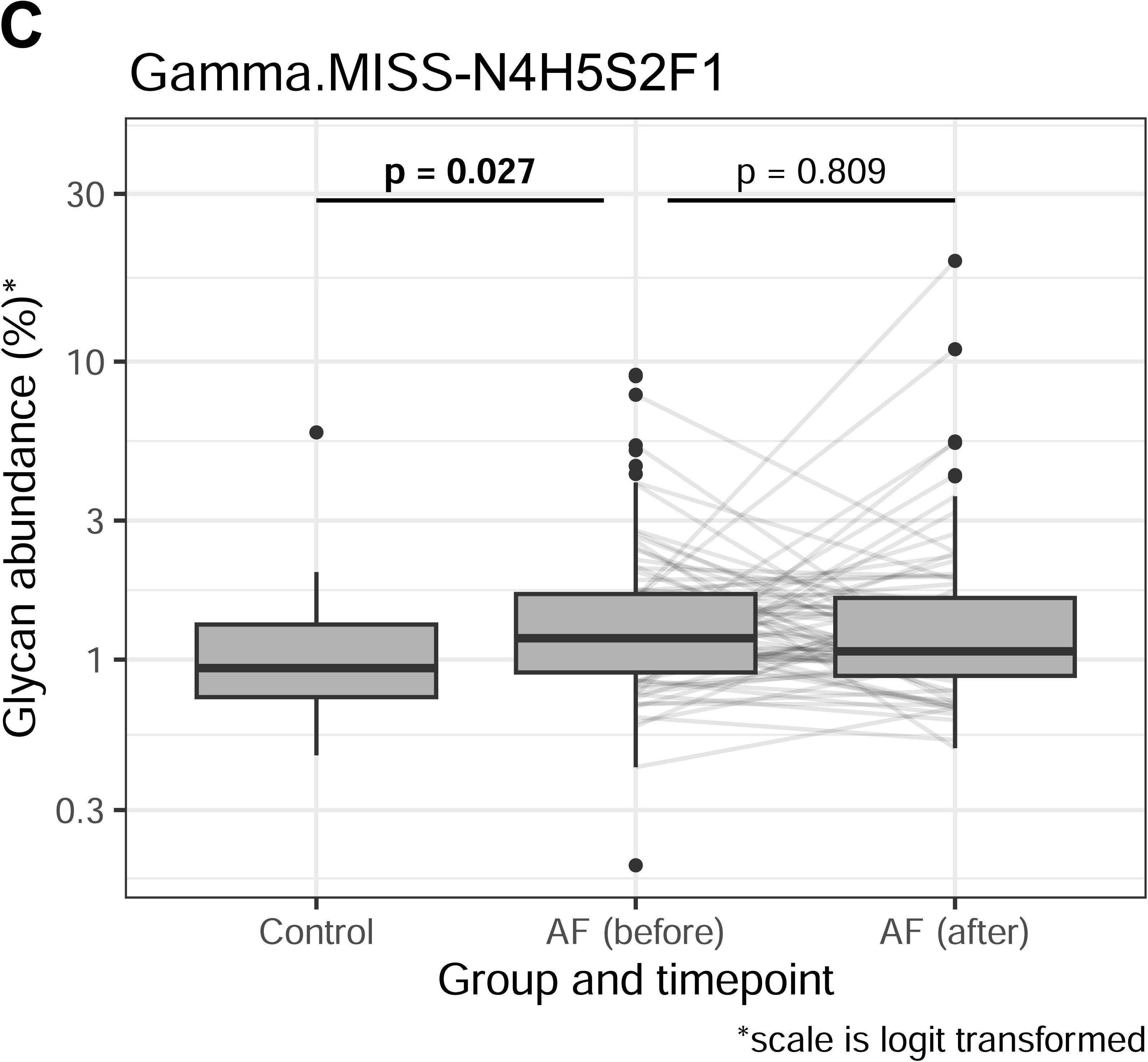
Schematic workflow of the high-throughput, site-specific N-glycosylation analysis method for human fibrinogen.

N-glycosylation profiling of human fibrinogen was initially performed on a plasma standard created from pooled plasma of the AF population. Glycosylation sites were identified based on the m/z values calculated from the masses of tryptic peptides carrying N-glycosylation sites and previously reported glycan structures (21,22). LC-MS analysis confirmed that the α chain was non-glycosylated, while both the β and γ chains displayed similar oligosaccharide structures. Glycopeptides with missed cleavages were observed at both β and γ glycosylation sites. Table 1 provides the glycan compositions of each site, along with abbreviations for naming fibrinogen glycoforms which passed the QC criteria (see Data processing in the Experimental procedures section for more details about the QC criteria). Variations in sample preparation – such as additional denaturation or increased concentrations of dithiothreitol, iodoacetamide, and trypsin – did not significantly reduce the occurrence of glycopeptides with missed cleavages. Glycopeptides with missed cleavages (Beta.MISS and Gamma.MISS) co-eluted with their fully digested counterparts (Beta.normal and Gamma.normal) with nearly identical retention times. A typical chromatogram with extracted ion traces of the most abundant glycopeptides from each glycosylation site is shown in Figure 2 and an example summed MS spectrum for both Beta and Gamma glycosylation sites is shown in Figure 3. The intensity of Beta.MISS glycopeptides was negligible compared to that of Beta.normal glycopeptides, failing to meet QC criteria in some samples. In contrast, Gamma.MISS glycopeptides were predominant over Gamma.normal glycopeptides and demonstrated significantly better repeatability (see below, Table S3). Consequently, Beta.normal and Gamma.MISS glycopeptides were used in the analyses.

**Figure 2.**
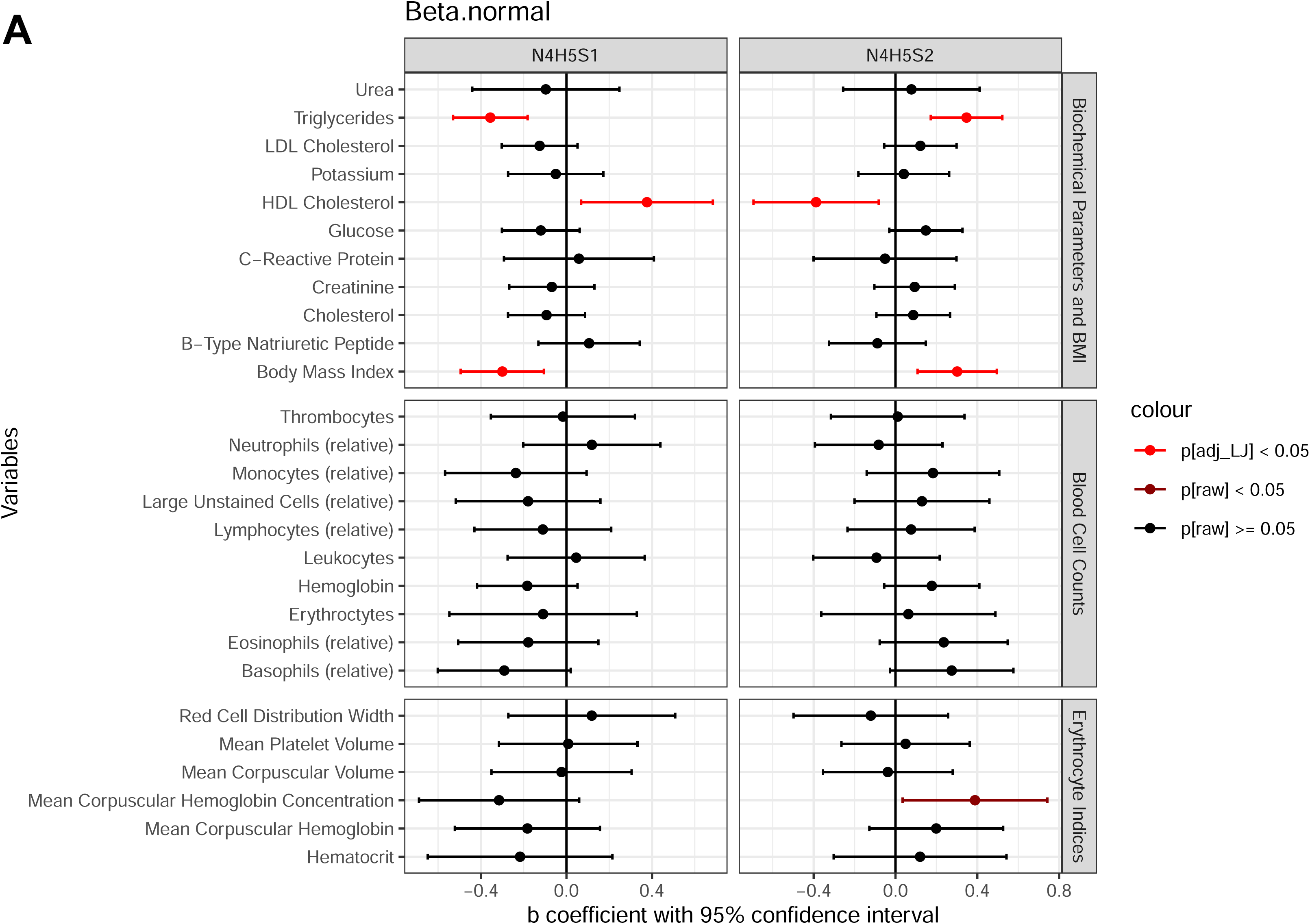
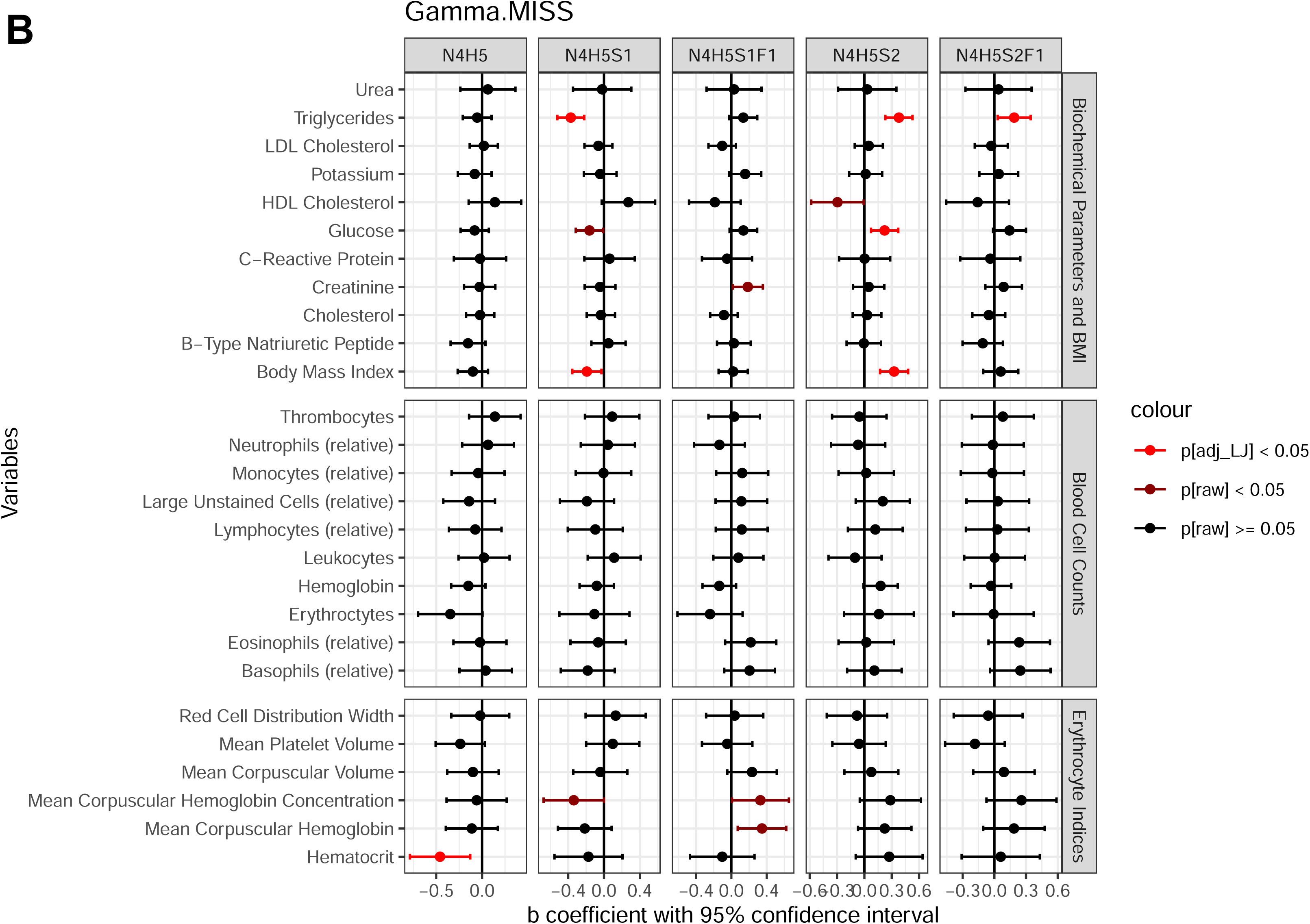
LC-MS analysis of fibrinogen N-glycosylation using the newly developed method: representative chromatogram with extracted ion traces of major glycoforms from two N-glycosylation sites, including detected missed cleavages.

**Figure 3.**
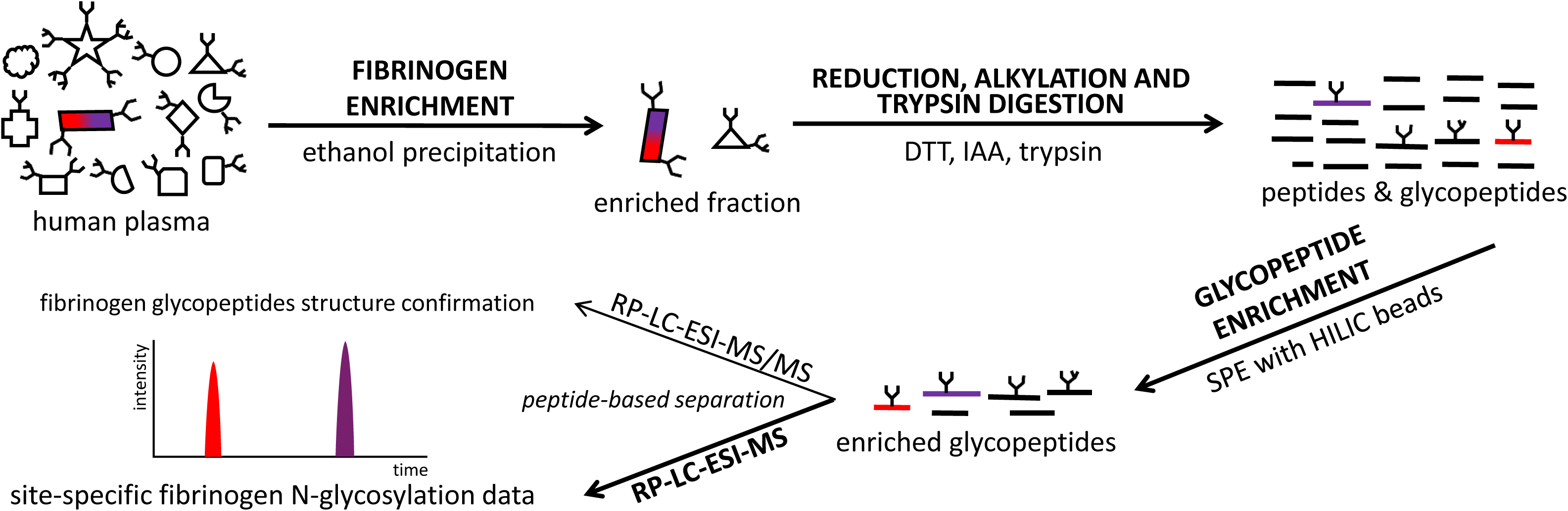
Typical summed mass spectra. *A*, Beta glycosylation site peak cluster with annotated triply [M+3H]^3+^ charged glycopeptide ions. *B*, Gamma glycosylation site peak cluster with annotated triply [M+3H]^3+^ charged glycopeptide ions.

**Table 1.**
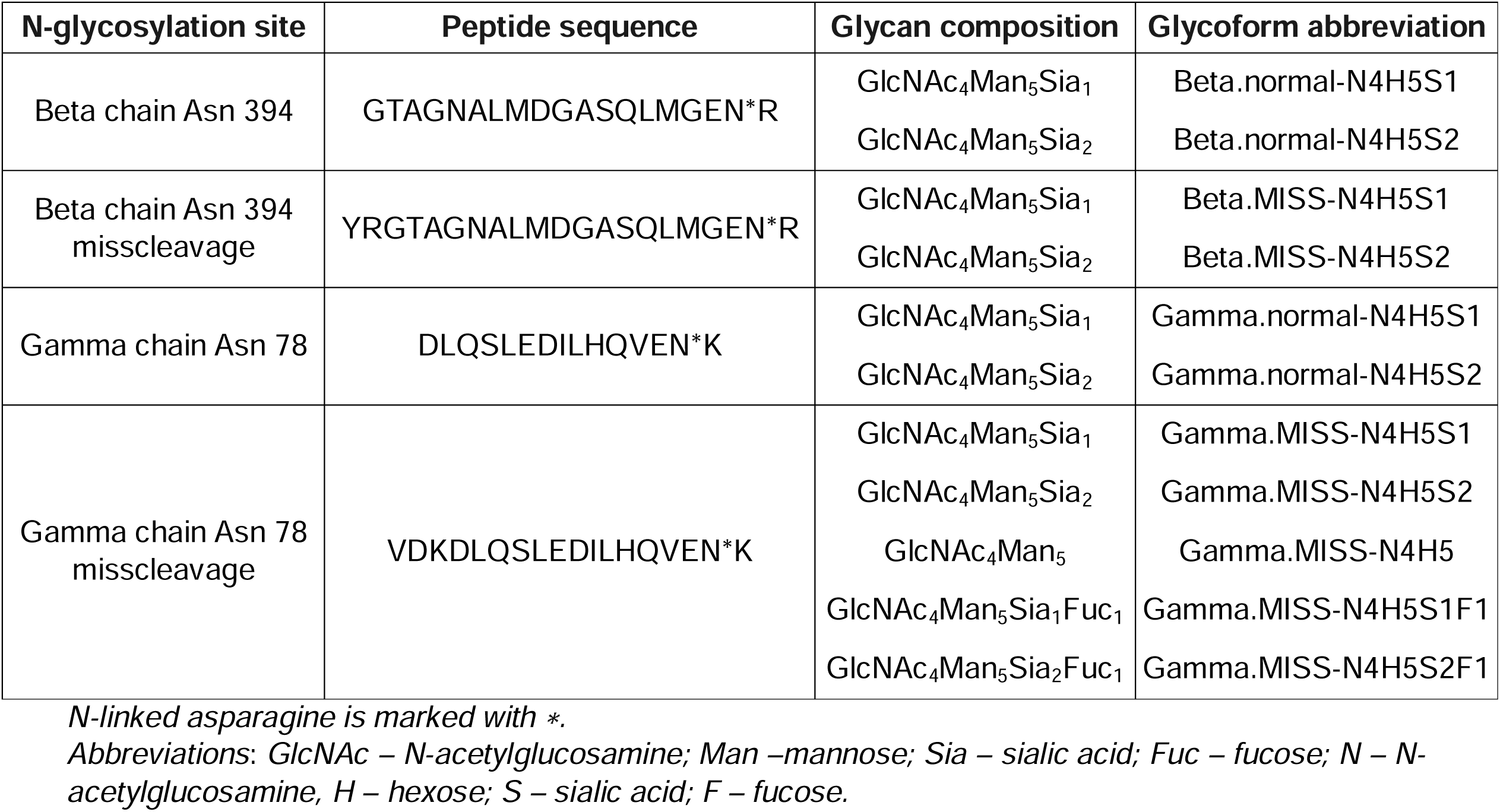
Trypsin digested peptide sequence for each fibrinogen glycosylation site with corresponding glycan compositions and glycoform abbreviations.

| N-glycosylation site | Peptide sequence | Glycan composition | Glycoform abbreviation |
| --- | --- | --- | --- |
| Beta chain Asn 394 | GTAGNALMDGASQLMGEN*R | GlcNAc<sub>4</sub>Man<sub>5</sub>Sia<sub>1</sub> | Beta.normal-N4H5S1 |
|  |  | GlcNAc<sub>4</sub>Man<sub>5</sub>Sia<sub>2</sub> | Beta.normal-N4H5S2 |
| Beta chain Asn 394<br>misscleavage | YRGTAGNALMDGASQLMGEN*R | GlcNAc<sub>4</sub>Man<sub>5</sub>Sia<sub>1</sub> | Beta.MISS-N4H5S1 |
|  |  | GlcNAc<sub>4</sub>Man<sub>5</sub>Sia<sub>2</sub> | Beta.MISS-N4H5S2 |
| Gamma chain Asn 78 | DLQSLEDILHQVEN*K | GlcNAc<sub>4</sub>Man<sub>5</sub>Sia<sub>1</sub> | Gamma.normal-N4H5S1 |
|  |  | GlcNAc<sub>4</sub>Man<sub>5</sub>Sia<sub>2</sub> | Gamma.normal-N4H5S2 |
| Gamma chain Asn 78<br>misscleavage | VDKDLQSLEDILHQVEN*K | GlcNAc<sub>4</sub>Man<sub>5</sub>Sia<sub>1</sub> | Gamma.MISS-N4H5S1 |
|  |  | GlcNAc<sub>4</sub>Man<sub>5</sub>Sia<sub>2</sub> | Gamma.MISS-N4H5S2 |
|  |  | GlcNAc<sub>4</sub>Man<sub>5</sub> | Gamma.MISS-N4H5 |
|  |  | GlcNAc<sub>4</sub>Man<sub>5</sub>Sia<sub>1</sub>Fuc<sub>1</sub> | Gamma.MISS-N4H5S1F1 |
|  |  | GlcNAc<sub>4</sub>Man<sub>5</sub>Sia<sub>2</sub>Fuc<sub>1</sub> | Gamma.MISS-N4H5S2F1 |

*N-linked asparagine is marked with \*.*
*Abbreviations: GlcNAc – N-acetylglucosamine; Man – mannose; Sia – sialic acid; Fuc – fucose; N – N-acetylglucosamine, H – hexose; S – sialic acid; F – fucose.*

All glycoforms included in the analyses were confirmed by MS/MS, as illustrated by the typical fragmentation spectra for Beta.normal-N4H5S1 and Gamma.MISS-N4H5S1 presented in Figure 4. Other major glycoforms (Beta.normal-N4H5S2 and Gamma.MISS-N4H5S2) displayed consistent fragmentation patterns, with MS/MS spectra provided in supplemental Figure S2. MS/MS spectra for less abundant glycoforms (Gamma.MISS-N4H5, Gamma.MISS-N4H5S1F1 and Gamma.MISS-N4H5S2F1) were of lower intensity but still yielded partial structural information (Figure S3).

**Figure 4.**
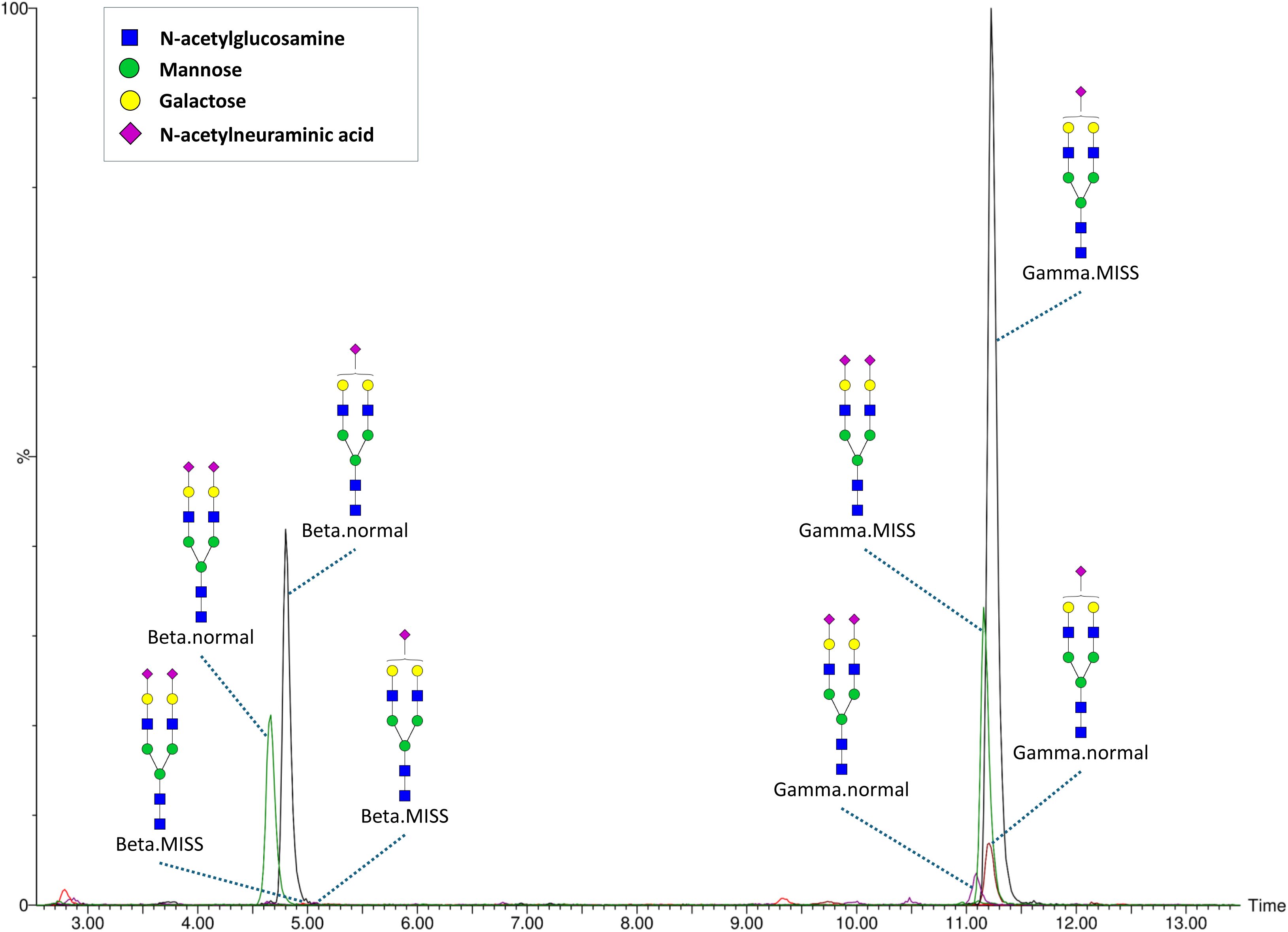
MS/MS fragmentation spectra of major glycoforms from both fibrinogen glycosylation sites. *A*, fragmentation pattern of Beta.normal-N4H5S1 [M+3H]^3+^ glycopeptide. *B*, fragmentation pattern of Gamma.MISS-N4H5S1 [M+3H]^3+^ glycopeptide.

Furthermore, we analyzed two series of pooled plasma standard replicates, which tested intra-plate (8 replicates were randomized across 1 plate) and inter-plate (16 replicates were randomized across 4 different plates) repeatability (Table S3). Average CV was 5% (SD = 4%) for the intra-plate and 12% (SD = 12%) for the inter-plate replicates. As anticipated, the major glycoforms (Beta.normal-N4H5S1, Beta.normal-N4H5S2 and Gamma.MISS-N4H5S1, Gamma.MISS-N4H5S2) exhibited significantly lower CVs compared to the less abundant glycoforms. Repeatability results suggest that the method is suitable for high-throughput profiling, although, like other high-throughput glycosylation analysis methods, it is subject to batch effects. Therefore, batch effect correction was applied to all analyzed samples (see Experimental Procedures for details).

### Intra-individual temporal stability study

An important question in glycosylation research is the temporal stability of glycan profiles within individuals and their variability between individuals. This aspect has not yet been examined for fibrinogen. To explore this, we conducted a small-scale study in which plasma samples from 14 healthy, age- and sex-matched individuals were analyzed at three time points. Samples were collected with intervals of 6 and 4 weeks, respectively, which – given fibrinogen’s approximate 4-day half-life in plasma (53) – should reflect predominantly newly synthesized protein at each time point.

The data from this experiment was processed to calculate two categories of coefficients of variation (CV) for each glycopeptide: intra-individual CV, calculated from the longitudinal samples of each subject, and inter-individual CV, derived from comparisons of samples across subjects at each time point. Six glycopeptides met the QC criteria (Gamma.MISS-N4H5S2F1 did not pass). The data (Table 2) indicate that inter-individual variation consistently exceeded intra-individual variation for all glycopeptides. Additionally, intra-individual variation was slightly higher than the method’s intra-plate repeatability.

**Table 2.** Temporal stability of fibrinogen N-glycosylation in healthy individuals over 10 weeks.

| **Glycopeptide** | **Intra-individual CV (%)** | **Inter-individual CV (%)** | **Method repeatability CV (%)** |
| --- | --- | --- | --- |
| Beta.normal-N4H5S1 | 1.8 | 3.8 | 1.1 |
| Beta.normal-N4H5S2 | 2.5 | 5.2 | 1.3 |
| Gamma.MISS-N4H5 | 11.1 | 18.7 | 12.5 |
| Gamma.MISS-N4H5S1 | 3.0 | 4.3 | 1.0 |
| Gamma.MISS-N4H5S1F1 | 13.7 | 23.2 | 4.7 |
| Gamma.MISS-N4H5S2 | 5.4 | 8.3 | 2.2 |

This stability within individuals and higher variability between individuals suggest that fibrinogen glycosylation patterns are consistent in healthy individuals, making them a potential biomarker candidate. Because the study subjects were age- and sex-matched, we observed inter-individual differences under conditions where these differences are expected to be minimal, suggesting that additional demographic or environmental factors could further amplify the observed variations.

Previous studies have similarly confirmed intra-individual temporal stability in N-glycosylation profiles of total plasma proteins, immunoglobulin G, alpha-1-acid glycoprotein, and complement component 3 (32,33,54,55). These findings reinforce that glycan profiles remain stable under physiological conditions but can shift in certain pathophysiological states, underscoring the diagnostic potential of glycosylation analysis.

### Fibrinogen N-glycome in atrial fibrillation

The diagnostic potential of the fibrinogen N-glycan profile was assessed in an AF population undergoing catheter ablation, with comparisons made to profiles from healthy controls.

Fibrinogen glycopeptides were associated with both sex and age (Table S4); therefore, these variables were included as covariates in all subsequent models. Regression analysis identified three glycopeptides significantly associated with AF, all of which were lower-abundance glycopeptides from the Gamma glycosylation site (Table 3). Specifically, Gamma.MISS-N4H5, Gamma.MISS-N4H5S1F1, and Gamma.MISS-N4H5S2F1 were found to be more abundant in AF patients compared to controls. The most notable finding was an increase in Gamma.MISS-N4H5, an asialylated glycoform. Asialylated fibrinogen is known to enhance clot formation, producing fibrin bundles with greater thickness compared to those formed by fibrinogen with disialylated glycans (24). Given that AF patients are predisposed to clot formation and routinely prescribed anticoagulants (45), these glycosylation changes could have functional implications in the prothrombotic state associated with AF. This underscores the potential of fibrinogen glycosylation alterations as both a mechanistic factor and a therapeutic target in AF management. However, further studies are needed to validate these findings. Statistically significant associations between AF and fibrinogen glycoformes are illustrated in Figure 5, along with data from 108 individuals who were resampled six months later. The fibrinogen N-glycan profile in these individuals remained unchanged over this period (Table 3), indicating that the distinct AF-associated glycan profile is stable over at least six months following the ablation procedure. It is worth noting that a previous analysis of the total plasma glycome, which includes fibrinogen, was performed on the same set of samples. The only statistically significant difference observed between AF patients and healthy controls was in the oligomannose structure Man9 (46), which is not present in fibrinogen. This underscores the importance of single-protein glycosylation analysis for biomarker discovery, as it provides a more precise and targeted approach than total plasma glycome profiling.

**Figure 5.**
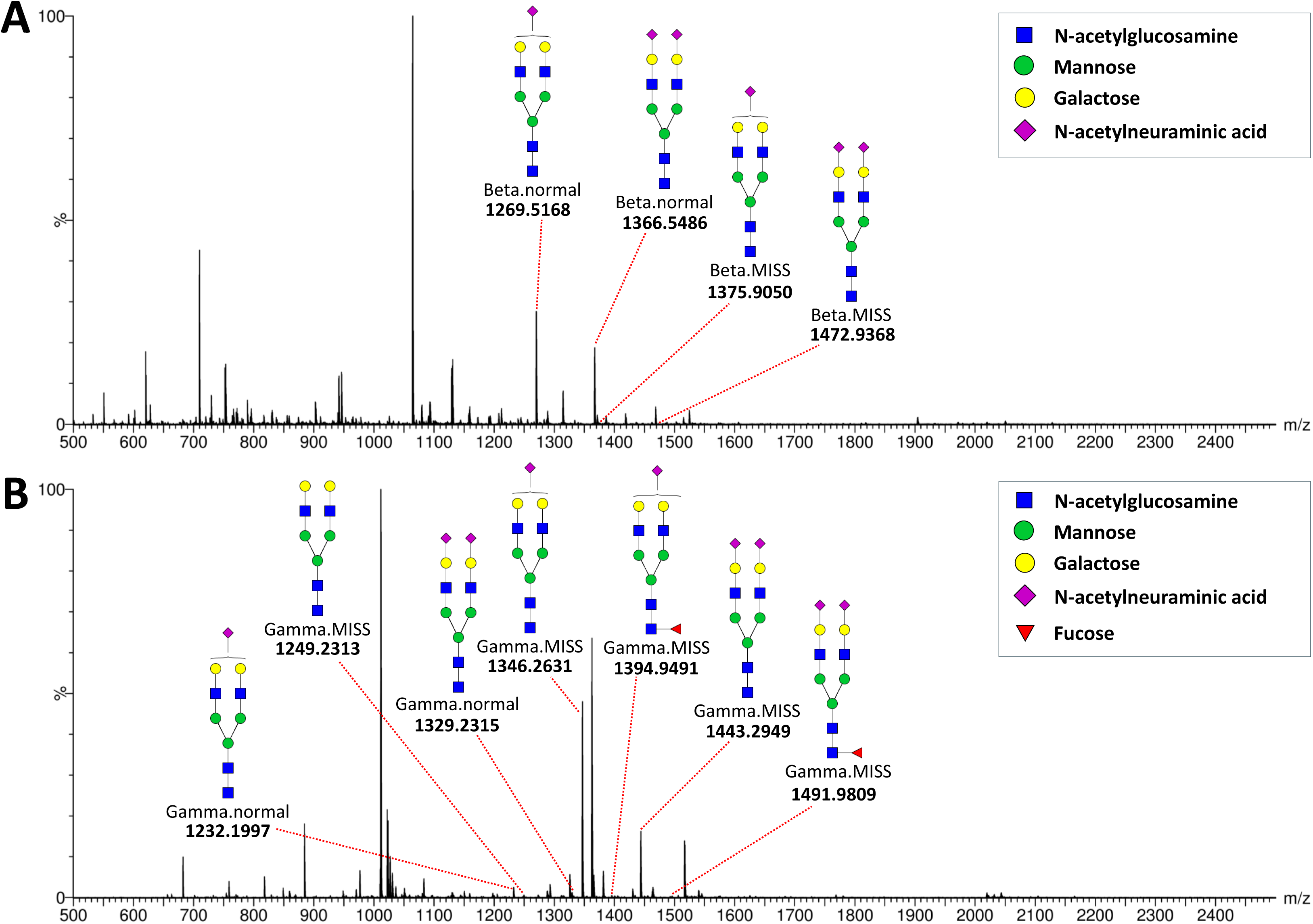
Fibrinogen glycopeptides significantly associated with atrial fibrillation, along with glycosylation data from 108 patients who were resampled six months later. *A*, Gamma.MISS-N4H5. *B*, Gamma.MISS-N4H5S1F1. *C*, Gamma.MISS-N4H5S2F1.

**Table 3.**
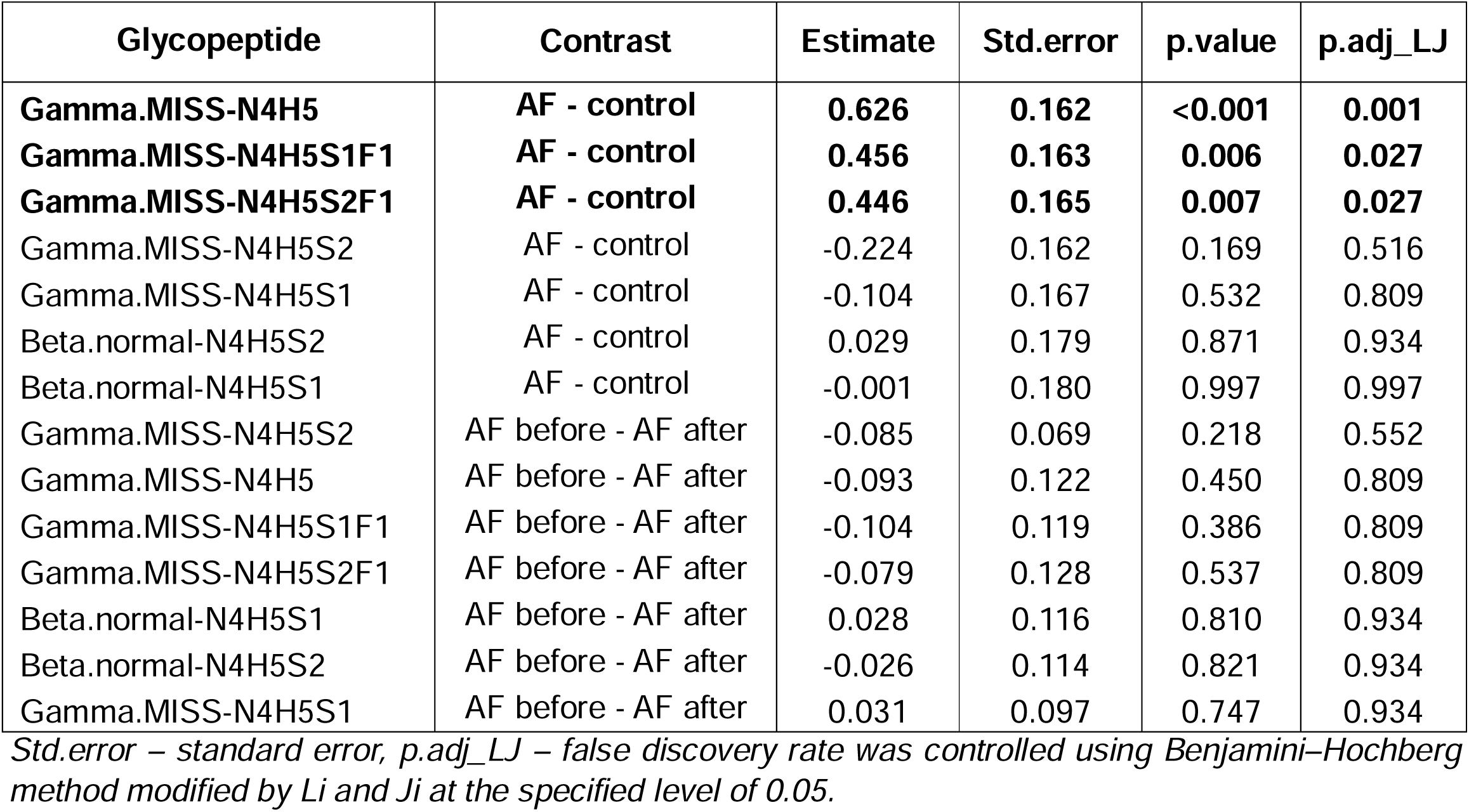
Differences in fibriongen N-glycosylation between AF patients and healthy controls (AF – control) and differences in fibrinogen N-glycosylation before vs. six months after the catheter ablation procedure in 108 individuals with AF (AF before – AF after). Statistically significant glycopeptides are highlighted in bold.

| Glycopeptide | Contrast | Estimate | Std.error | p.value | p.adj_LJ |
| --- | --- | --- | --- | --- | --- |
| **Gamma.MISS-N4H5** | **AF - control** | **0.626** | **0.162** | **<0.001** | **0.001** |
| **Gamma.MISS-N4H5S1F1** | **AF - control** | **0.456** | **0.163** | **0.006** | **0.027** |
| **Gamma.MISS-N4H5S2F1** | **AF - control** | **0.446** | **0.165** | **0.007** | **0.027** |
| Gamma.MISS-N4H5S2 | AF - control | -0.224 | 0.162 | 0.169 | 0.516 |
| Gamma.MISS-N4H5S1 | AF - control | -0.104 | 0.167 | 0.532 | 0.809 |
| Beta.normal-N4H5S2 | AF - control | 0.029 | 0.179 | 0.871 | 0.934 |
| Beta.normal-N4H5S1 | AF - control | -0.001 | 0.180 | 0.997 | 0.997 |
| Gamma.MISS-N4H5S2 | AF before - AF after | -0.085 | 0.069 | 0.218 | 0.552 |
| Gamma.MISS-N4H5 | AF before - AF after | -0.093 | 0.122 | 0.450 | 0.809 |
| Gamma.MISS-N4H5S1F1 | AF before - AF after | -0.104 | 0.119 | 0.386 | 0.809 |
| Gamma.MISS-N4H5S2F1 | AF before - AF after | -0.079 | 0.128 | 0.537 | 0.809 |
| Beta.normal-N4H5S1 | AF before - AF after | 0.028 | 0.116 | 0.810 | 0.934 |
| Beta.normal-N4H5S2 | AF before - AF after | -0.026 | 0.114 | 0.821 | 0.934 |
| Gamma.MISS-N4H5S1 | AF before - AF after | 0.031 | 0.097 | 0.747 | 0.934 |

*Std.error – standard error, p.adj\_LJ – false discovery rate was controlled using Benjamini–Hochberg method modified by Li and Ji at the specified level of 0.05.*

We also examined potential differences in fibrinogen glycosylation between patients who experienced AF recurrence within six months post-catheter ablation and those who did not. To assess this, we tested several predictive models: glycan profiles of patients with AF recurrence versus those without at the initial time point, profiles of recurrence versus non-recurrence at the six-month follow-up, and changes in glycan profiles over time (delta) between the two time points for patients with and without recurrence. Results from these regression models are provided in Table S5, where no significant associations were found between the fibrinogen N-glycan profile and AF recurrence.

Leveraging the available clinical data for all study participants, we conducted the first investigation into potential associations between fibrinogen N-glycosylation and a range of biochemical, hematological parameters, as well as BMI. Results from the regression model, adjusted for age and sex, are presented in Figure 6 and Table S6. We identified strong associations between triglyceride levels and several fibrinogen glycoforms, including the major glycoforms at both glycosylation sites: Beta.normal-N4H5S1, Beta.normal-N4H5S2, Gamma.MISS-N4H5S1, and Gamma.MISS-N4H5S2, as well as a minor glycoform at the Gamma site, Gamma.MISS-N4H5S2F1. Mono-sialylated glycoforms were negatively associated with triglycerides, whereas di-sialylated glycoforms showed positive associations. Similar trends were observed for BMI, with the major glycoforms (Beta.normal-N4H5S1, Beta.normal-N4H5S2, Gamma.MISS-N4H5S1, and Gamma.MISS-N4H5S2) showing consistent patterns with those seen for triglycerides. In contrast, significant associations were observed with HDL cholesterol but in opposite directions, and only at the Beta glycosylation site – Beta.normal-N4H5S1 was positively associated with HDL, while Beta.normal-N4H5S2 was negatively associated. Additionally, Gamma.MISS-N4H5S2 showed a significant positive association with glucose levels, and Gamma.MISS-N4H5 exhibited a significant negative association with hematocrit. Elevated triglycerides, BMI, and glucose levels are well-established risk factors for cardiovascular diseases, and in this study, they were associated with increased fibrinogen sialylation. Plasma triglyceride levels have been shown to influence fibrinogen clearance, extending its half-life (56). Additionally, asialofibrinogen, along with other asialylated plasma proteins, is removed from circulation via the asialoglycoprotein receptor, and no circulating fibrinogen molecules completely lacking both sialic acid residues have been detected (57). Given that elevated plasma fibrinogen levels are common in patients with cardiovascular diseases (15), the observed link between increased sialylation and these risk factors is not unexpected. These findings underscore the intricate relationship between fibrinogen glycosylation and cardiovascular risk factors, suggesting that changes in sialylation may reflect a compensatory mechanism or a pathophysiological response. Further studies are needed to explore whether targeting fibrinogen glycosylation could offer new insights into disease mechanisms or potential therapeutic opportunities.

**Figure 6.**
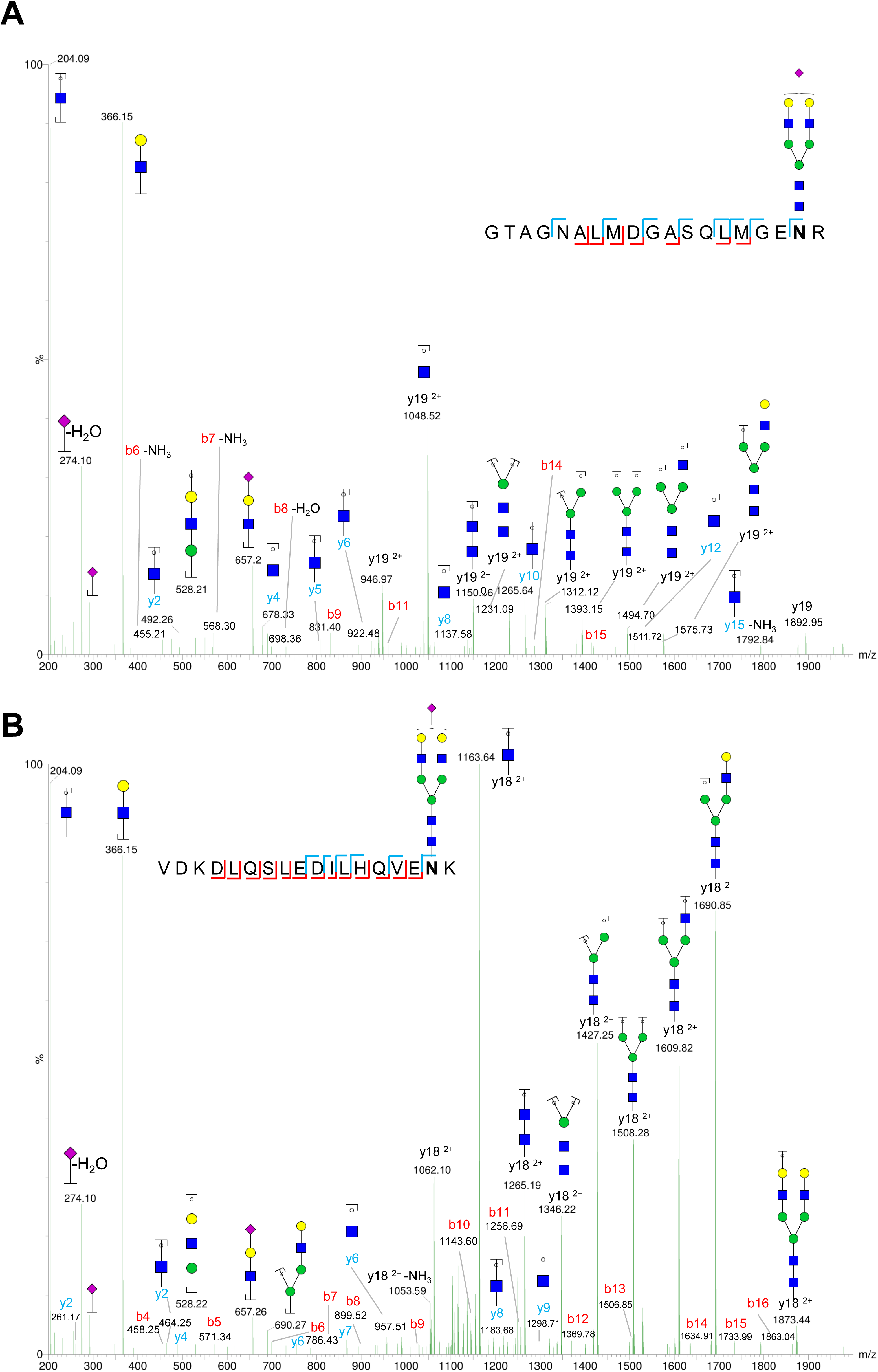
Associations between fibrinogen glycoforms and range of biochemical, hematological parameters, as well as BMI. *A*, effects of prediction models for Beta.normal glycopeptides. *B*, effects of prediction models for Gamma.MISS glycopeptides.

## Conclusion

In this study, we developed and validated a high-throughput, site-specific method for the analysis of fibrinogen N-glycosylation. The method successfully identified key glycoforms associated with atrial fibrillation (AF) and established a stable N-glycosylation profile in healthy individuals over time. Notably, an increase in the asialylated glycoform Gamma.MISS-N4H5 was observed in AF patients, which may contribute to the prothrombotic state seen in this condition. The stability of fibrinogen glycosylation profiles in AF patients over a two-month period further supports their potential as biomarkers for AF. Additionally, associations between fibrinogen glycoforms and key biochemical and hematological parameters, such as triglycerides, BMI, and glucose, highlight the intricate relationship between fibrinogen glycosylation and cardiovascular risk factors. These findings not only establish fibrinogen glycosylation as a promising biomarker candidate for cardiovascular conditions but also provide a practical analytical pipeline for investigating protein-specific glycosylation in clinical samples, opening new avenues for biomarker discovery and mechanistic studies in cardiovascular diseases.

## Acknowledgments

This research was supported by the project “GLYCARD: Glycosylation in Cardiovascular Diseases” (UIP-2019-04-5692), funded by the Croatian Science Foundation.

The graphical abstract was created with BioRender.com.

## Data Availability

All raw mass spectrometry data files are publicly available on the PRIDE repository (http://www.ebi.ac.uk/pride) under the identifier: PXD058737 (Token: cpvRuOjb4JO4).

## Author contributions

T. K., I. G., and O. G. designed research; D. Š., J. S.-N., A. B., L. Đ., D. R. and T. K. performed research; D. K., D. Š., T. K., and I. G. analyzed data; T. K., D. Š. and I. G. wrote the paper.

## Abbreviations

AF: Atrial fibrillation
BEH: Bridged ethylene hybrid
BMI: Body mass index
BPI: Base peak intensity
CID: Collision induced dissociation
CV: Coefficient of variation
DAD: Data-dependent acquisition
ESI: Electrospray ionization
ESRD: End-stage renal disease
HILIC: Hydrophilic interaction chromatography
MRM: Multiple reaction monitoring
QC: Quality control
RP: Reversed-phase
SPE: Solid-phase extraction
TFA: Trifluoroacetic acid
QTOF: Quadrupole time-of-flight
UPLC: Ultra-performance liquid chromatography

