## Supplementary material for "High-Throughput Site-Specific N-Glycosylation Profiling of Human Fibrinogen in Atrial Fibrillation": Figure S1

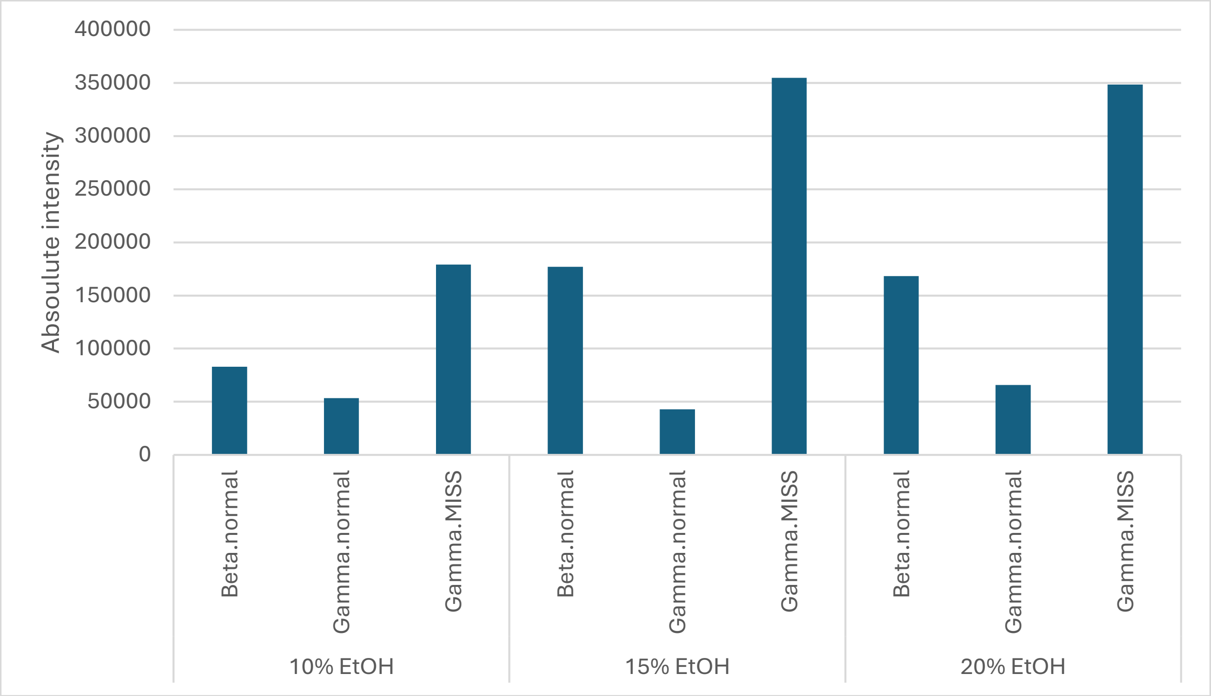


**Figure S1.** Absolute intensity of the most abundant fibrinogen glycopeptide peak, N4H5S1 $[M+3H]^{3+}$, across different peptides, measured from 20 μL of human plasma precipitated with 10%, 15%, and 20% absolute ethanol.
