## Supplementary material for "High-Throughput Site-Specific N-Glycosylation Profiling of Human Fibrinogen in Atrial Fibrillation": Figure S3

A

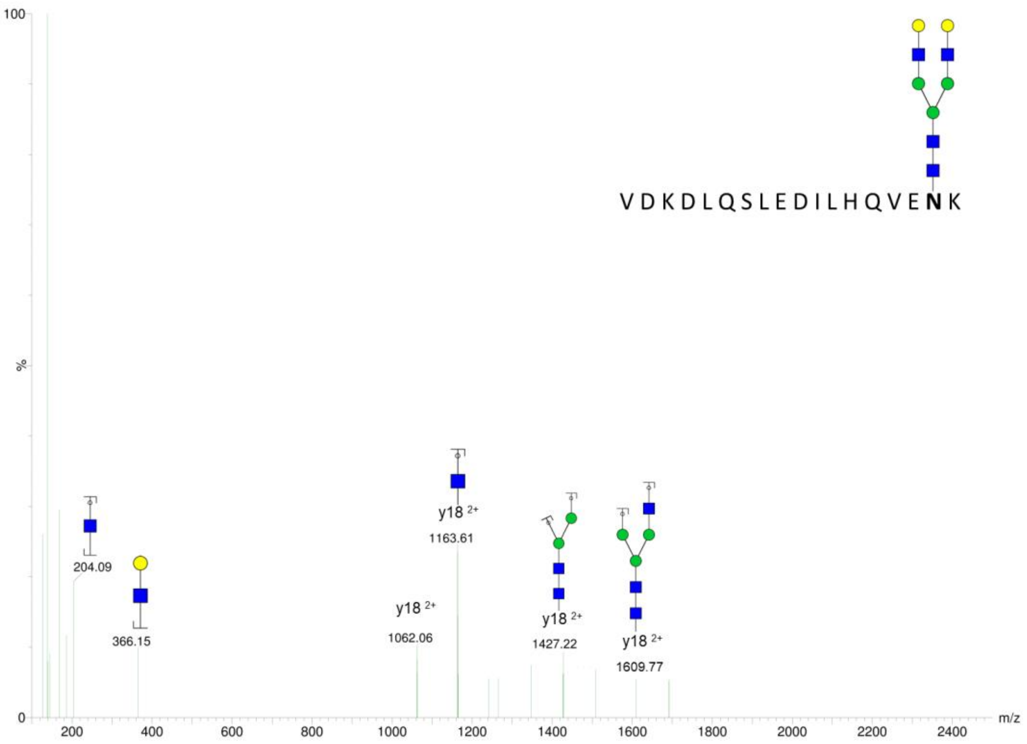
VDKDLQSLEDILHQVENK
100
%
0
204.09
366.15
y18 2+
1163.61
y18 2+
1062.06
y18 2+
1427.22
y18 2+
1609.77
200
400
600
800
1000
1200
1400
1600
1800
2000
2200
2400
m/z

B

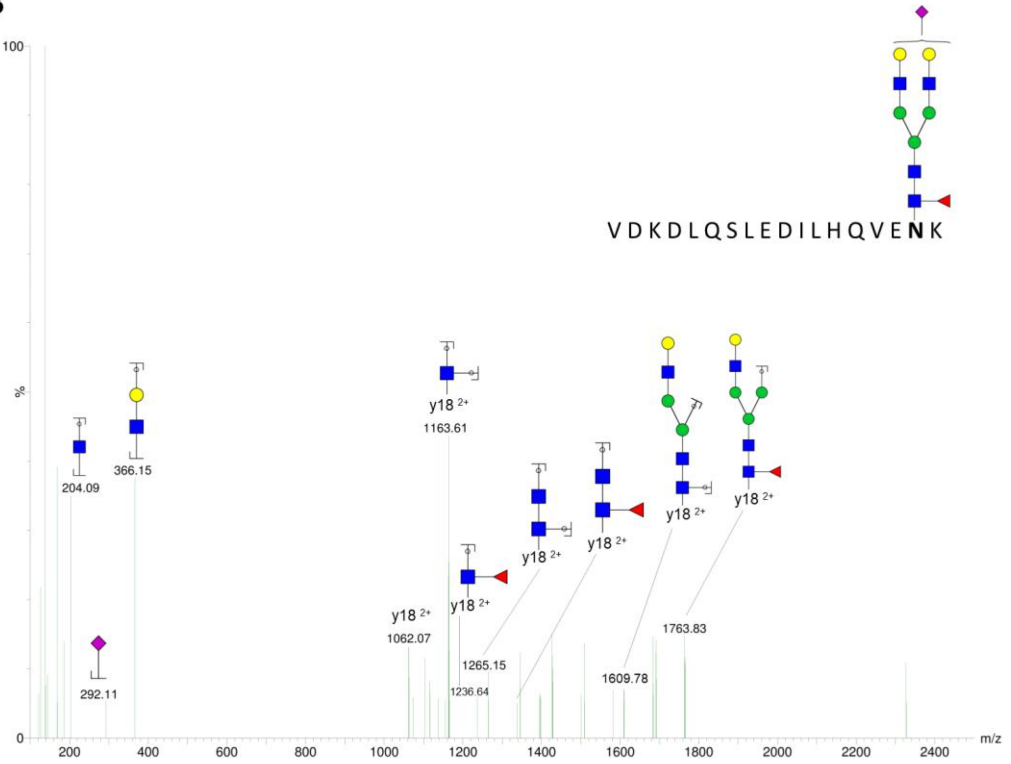
VDKDLQSLEDILHQVENK
100
%
0
204.09
366.15
292.11
y18 2+
1163.61
y18 2+
1062.07
y18 2+
1236.64
y18 2+
1265.15
y18 2+
y18 2+
1609.78
y18 2+
1763.83
200
400
600
800
1000
1200
1400
1600
1800
2000
2200
2400
m/z

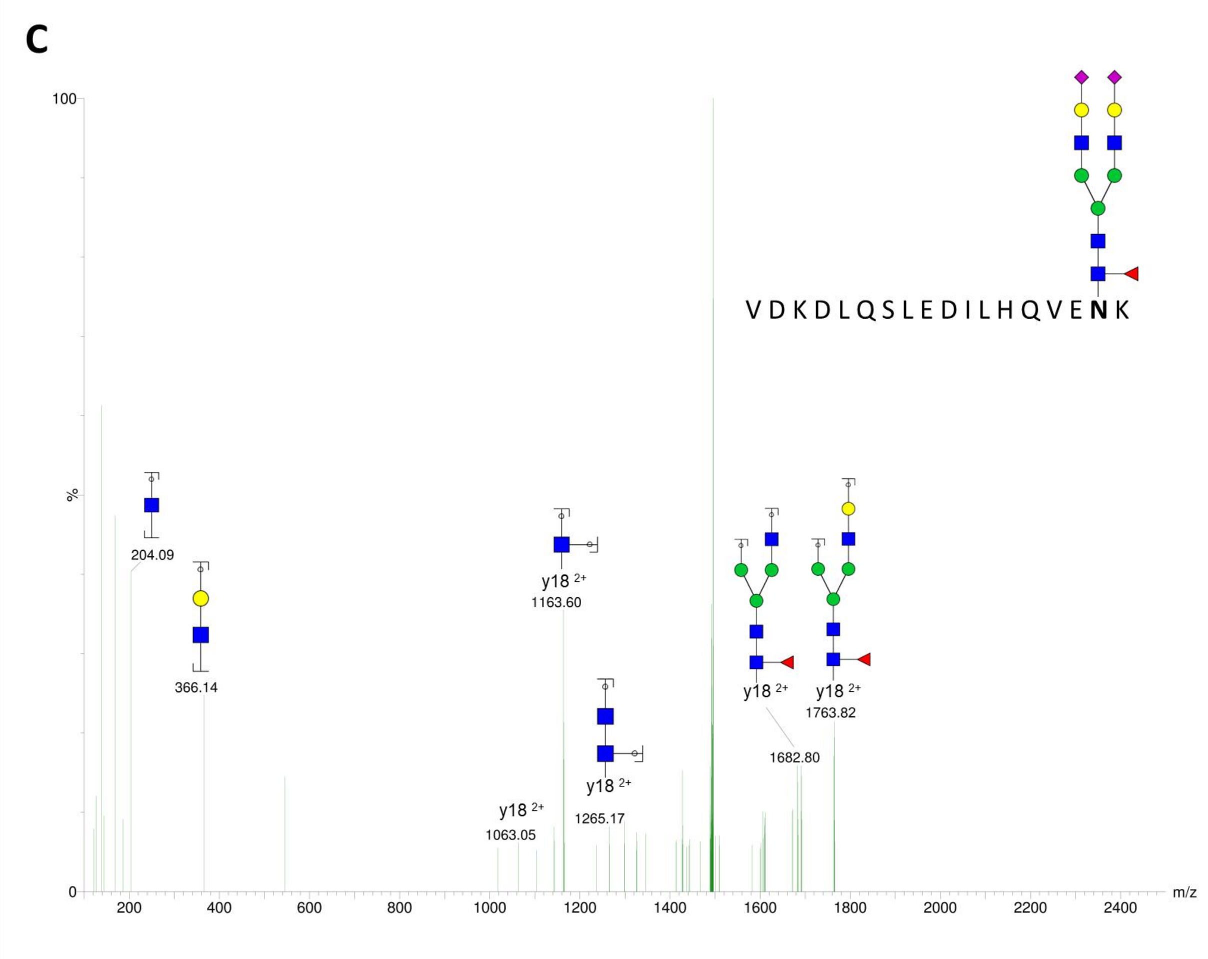


**Figure S3.** MS/MS fragmentation spectra of minor glycoforms from the fibrinogen Gamma glycosylation site**.** *A*, fragmentation pattern of Gamma.MISS-N4H5 $[M+3H]^{3+}$ glycopeptide. *B*, fragmentation pattern of Gamma.MISS-N4H5S1F1 $[M+3H]^{3+}$ glycopeptide. *C,* fragmentation pattern of Gamma.MISS-N4H5S2F1 $[M+3H]^{3+}$ glycopeptide.
