## Supplementary material for "High-Throughput Site-Specific N-Glycosylation Profiling of Human Fibrinogen in Atrial Fibrillation": Figure S2

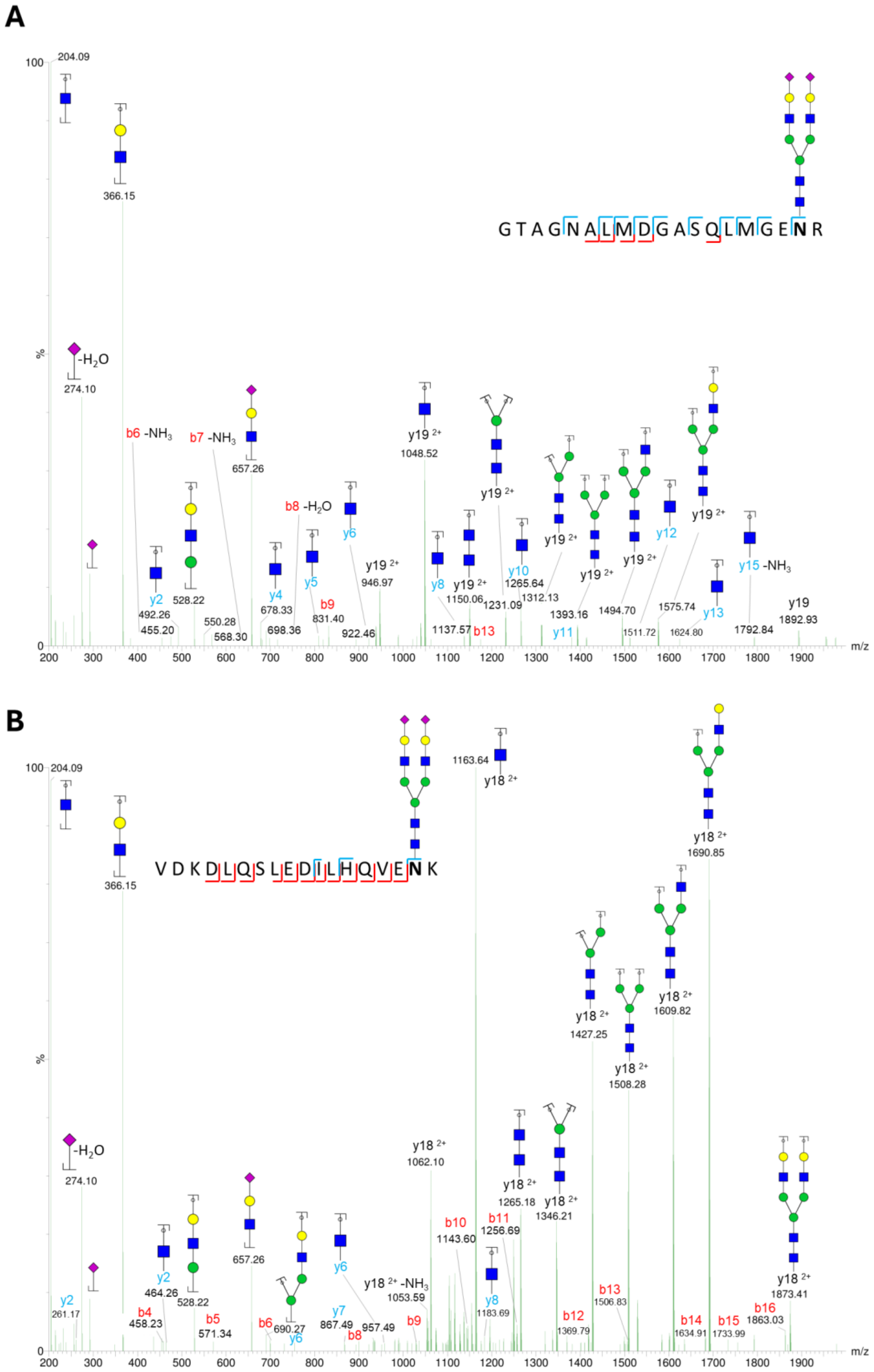


**Figure S2.** MS/MS fragmentation spectra of N4H5S2 glycoforms from both fibrinogen glycosylation sites**.** *A*, fragmentation pattern of Beta.normal-N4H5S2 $[M+3H]^{3+}$ glycopeptide. *B*, fragmentation pattern of Gamma.MISS-N4H5S2 $[M+3H]^{3+}$ glycopeptide.
