## Supplementary for "High-Throughput Site-Specific N-Glycosylation Profiling of Human Fibrinogen in Atrial Fibrillation": Supplementary Figures.pdf

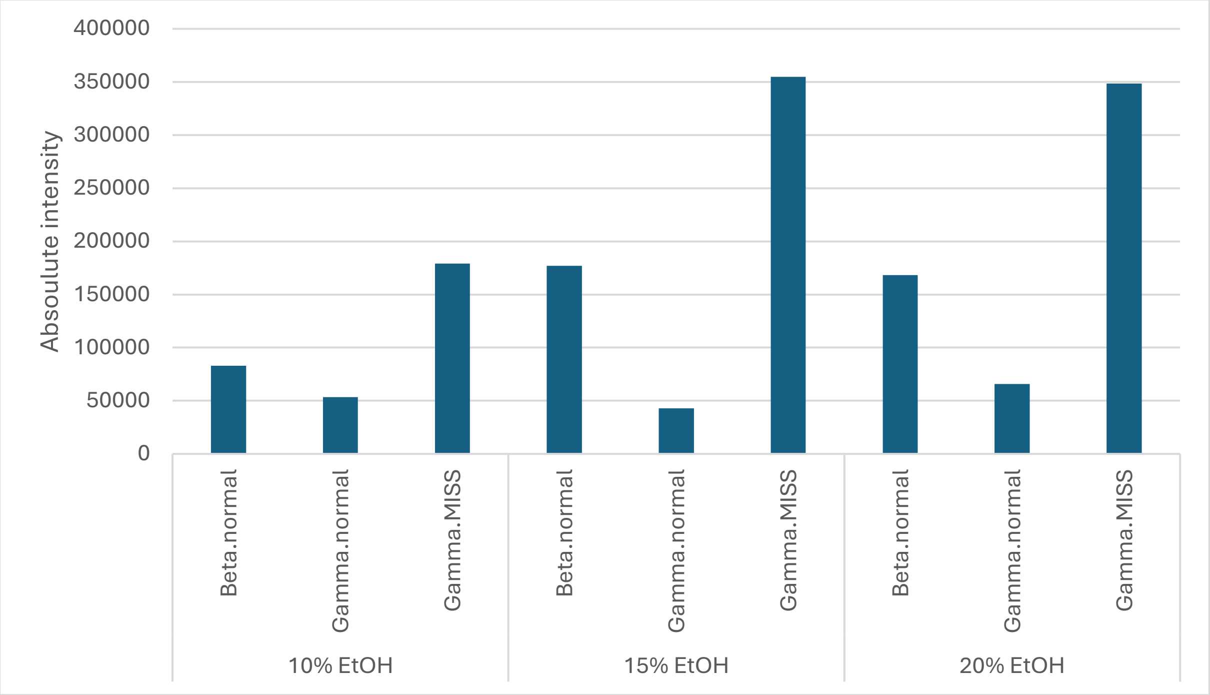


**Figure S1.** Absolute intensity of the most abundant fibrinogen glycopeptide peak, N4H5S1 $[M+3H]^{3+}$, across different peptides, measured from 20 µL of human plasma precipitated with 10%, 15%, and 20% absolute ethanol.

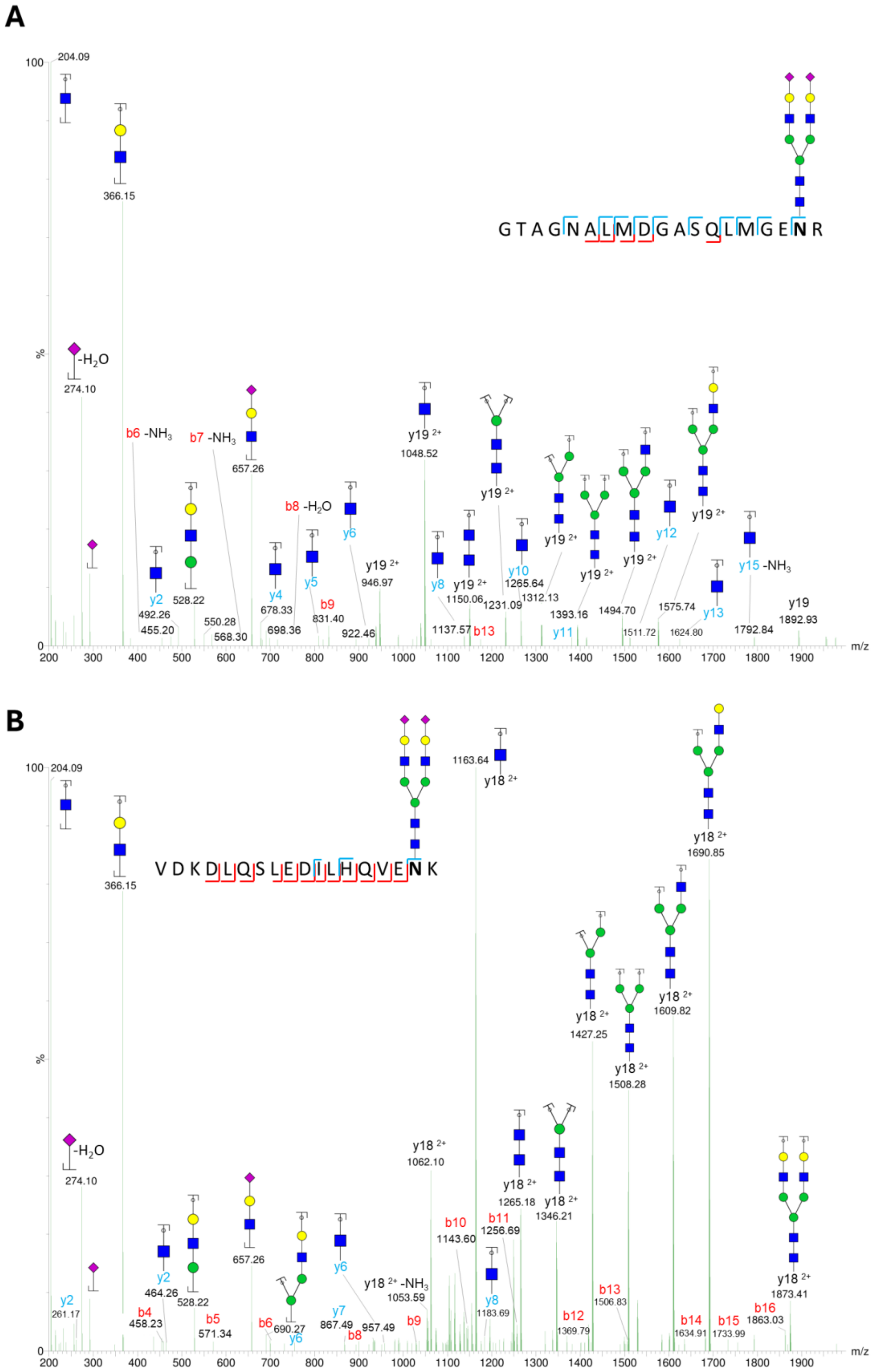


**Figure S2.** MS/MS fragmentation spectra of N4H5S2 glycoforms from both fibrinogen glycosylation sites**.** *A*, fragmentation pattern of Beta.normal-N4H5S2 $[M+3H]^{3+}$ glycopeptide. *B*, fragmentation pattern of Gamma.MISS-N4H5S2 $[M+3H]^{3+}$ glycopeptide.

**A**

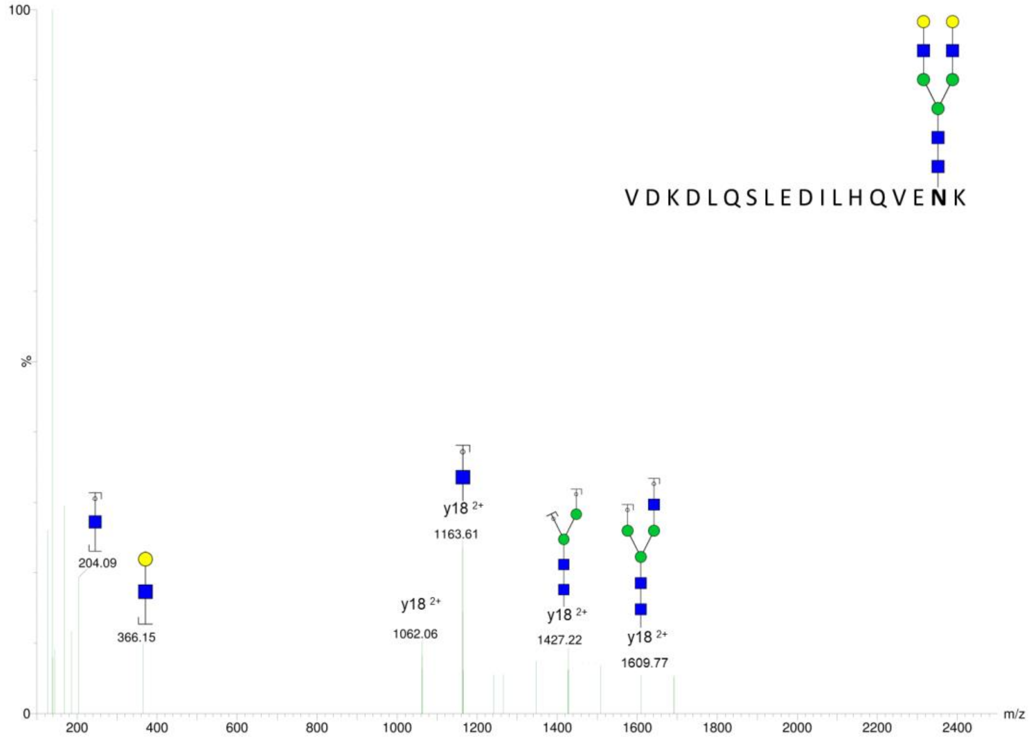
100
%
0
VDKDLQSLEDILHQVENK
204.09
366.15
y18 2+
1062.06
y18 2+
1163.61
y18 2+
1427.22
y18 2+
1609.77
200
400
600
800
1000
1200
1400
1600
1800
2000
2200
2400
m/z

**B**

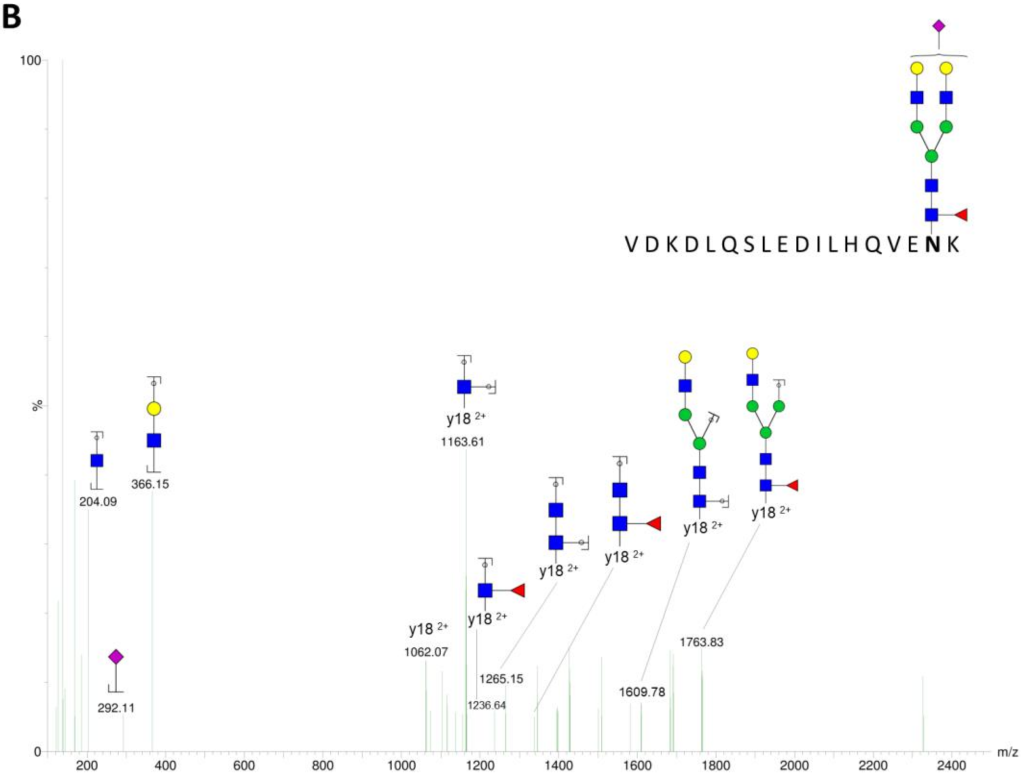
100
%
0
VDKDLQSLEDILHQVENK
204.09
366.15
292.11
y18 2+
1062.07
y18 2+
1163.61
y18 2+
1236.64
y18 2+
1265.15
y18 2+
y18 2+
1609.78
y18 2+
1763.83
200
400
600
800
1000
1200
1400
1600
1800
2000
2200
2400
m/z

C

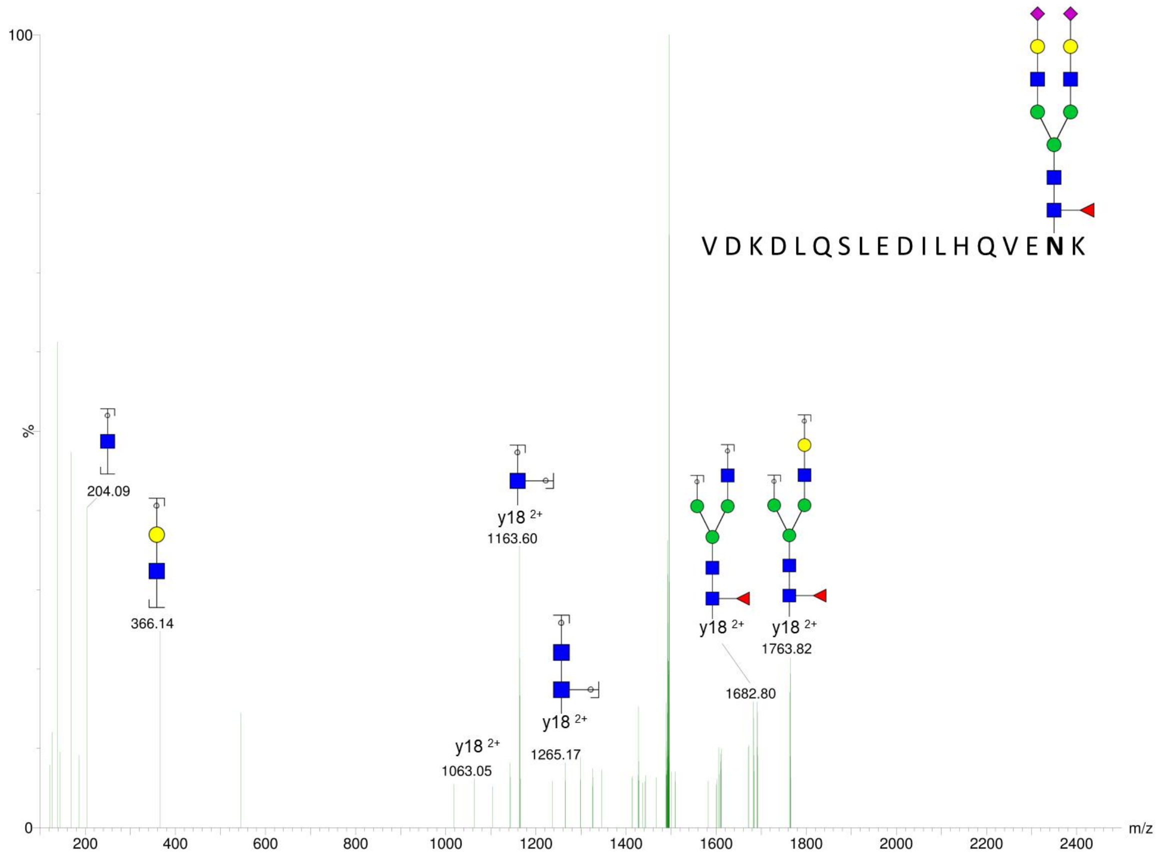


**Figure S3.** MS/MS fragmentation spectra of minor glycoforms from the fibrinogen Gamma glycosylation site**.** *A*, fragmentation pattern of Gamma.MISS-N4H5 $[M+3H]^{3+}$ glycopeptide. *B*, fragmentation pattern of Gamma.MISS-N4H5S1F1 $[M+3H]^{3+}$ glycopeptide. *C,* fragmentation pattern of Gamma.MISS-N4H5S2F1 $[M+3H]^{3+}$ glycopeptide.
