## Supplementary for "High-Throughput Site-Specific N-Glycosylation Profiling of Human Fibrinogen in Atrial Fibrillation": Table S1.pdf

**Table S1.** Characteristics of the study population. Age and BMI are given as the median and interquartile range. All other categorical variables are presented as the N (%). AF – atrial fibrillation, BMI – body mass index

| Variable | | AF | Healthy control |
|---|---|---|---|
| **N** | | 181 | 52 |
| **Age (years)** | | 63 (57-69) | 64 (57-69) |
| **Female sex** | | 62 (36%) | 18 (35%) |
| **BMI** | | 28.7 (26.1-31.6) | N/A |
| **Timepoint** | | | |
| | Before | 181 (100%) | 52 (100%) |
| | After | 108 (60%) | 0 (0%) |
| **AF recurrence after 6 months** | | | |
| | No | 68 (63%) | |
| | Yes | 34 (31%) | |
| | N/A | 6 (6%) | |
