## Supplementary for "High-Throughput Site-Specific N-Glycosylation Profiling of Human Fibrinogen in Atrial Fibrillation": Table S2.pdf

**Table S2.** Proteomic data analysis of the enriched fibrinogen glycopeptides following HILIC enrichment protocol. Sage was used as a proteomics database search engine. To analyze and visualize the results of the search, PeptideShaker was used. ProteomeID: UP000005640 9606.

| | Protein Inference | Main Accession | Description | Protein Group | Secondary Accessions | Chromosome | Gene Name | Taxonomy | Organism Identifier | Validated Coverage [%] | All Coverage [%] | Possible Coverage [%] | #Peptides | #Validated Peptides | #Unique Peptides | #Validated Unique Peptides | #PSMs | #Validated PSMs | Confidently Localized Modification Sites | #Confidently Localized Modification Sites | Ambiguously Localized Modification Sites | #Ambiguously Localized Modification Sites | Spectrum Counting | Relative abundance | MW [kDa] | Confidence [%] | Validation |
|---|---|---|---|---|---|---|---|---|---|---|---|---|---|---|---|---|---|---|---|---|---|---|---|---|---|---|---|
| 1 | Single Protein | P02768 | Albumin (ALBU_HUMAN) | P02768 | | 4 | ALB | Homo sapiens | 9606 | 24.14 | 24.14 | 86.54 | 9 | 9 | 9 | 9 | 296 | 294 | | | | | 15829.79471 | 0.450687757 | 69.32149566 | 100 | Confident |
| 2 | Single Protein | P01861 | Immunoglobulin heavy constant gamma 4 (IGHG4_HUMAN) | P01861 | | 14 | IGHG4 | Homo sapiens | 9606 | 5.81 | 5.81 | 59.09 | 1 | 1 | 1 | 1 | 7 | 7 | | | | | 848.8300584 | 0.024166916 | 43.80386763 | 100 | Doubtful |
| 3 | Single Protein | P02671 | Fibrinogen alpha chain (FIBA_HUMAN) | P02671 | | 4 | FGA | Homo sapiens | 9606 | 5.77 | 5.77 | 66.63 | 3 | 3 | 3 | 3 | 35 | 35 | | | | | 1721.197865 | 0.049003971 | 94.91441498 | 100 | Confident |
| 4 | Single Protein | P02679 | Fibrinogen gamma chain (FIBG_HUMAN) | P02679 | | 4 | FGG | Homo sapiens | 9606 | 27.37 | 27.37 | 90.51 | 6 | 6 | 6 | 6 | 78 | 75 | | | | | 5190.581019 | 0.147780269 | 51.47886685 | 100 | Confident |
| 5 | Single Protein | P02675 | Fibrinogen beta chain (FIBB_HUMAN) | P02675 | | 4 | FGB | Homo sapiens | 9606 | 20.16 | 20.16 | 81.06 | 5 | 5 | 5 | 5 | 66 | 66 | | | | | 4705.431235 | 0.133967641 | 55.89226078 | 100 | Confident |
| 6 | Single Protein | Q92845 | Kinesin-associated protein 3 (KIFA3_HUMAN) | Q92845 | | 1 | KIFAP3 | Homo sapiens | 9606 | 3.66 | 3.66 | 59.47 | 1 | 1 | 1 | 1 | 26 | 0 | | | | | 0 | 0 | 91.1464496 | 89.47368421 | Doubtful |
| 7 | Single Protein | P01857 | Immunoglobulin heavy constant gamma 1 (IGHG1_HUMAN) | P01857 | | 14 | IGHG1 | Homo sapiens | 9606 | 10.53 | 10.53 | 63.66 | 2 | 2 | 1 | 1 | 16 | 16 | | | | | 1787.412676 | 0.050889163 | 43.88410476 | 100 | Confident |
| 8 | Single Protein | O14917 | Protocadherin-17 (PCD17_HUMAN) | O14917 | | 13 | PCDH17 | Homo sapiens | 9606 | 1.29 | 1.29 | 68.94 | 1 | 1 | 1 | 1 | 1 | 1 | Oxidation of M (M1091); Oxidation of M (M1096) | Oxidation of M (2) | | | 35.513362 | 0.001011096 | 126.150807 | 100 | Doubtful |
| 9 | Single Protein | Q6GMV2 | Histone-lysine N-trimethyltransferase SMYD5 (SMYD5_HUMAN) | Q6GMV2 | | 2 | SMYD5 | Homo sapiens | 9606 | 4.55 | 4.55 | 57.66 | 1 | 1 | 1 | 1 | 1 | 1 | | | | | 117.7393205 | 0.003352139 | 47.30993361 | 100 | Doubtful |
| 10 | Single Protein | P01834 | Immunoglobulin kappa constant (IGKC_HUMAN) | P01834 | | 2 | IGKC | Homo sapiens | 9606 | 18.69 | 18.69 | 89.72 | 1 | 1 | 1 | 1 | 16 | 16 | | | | | 4729.196039 | 0.134644245 | 11.7577683 | 100 | Doubtful |
| 11 | Single Protein | P23471 | Receptor-type tyrosine-protein phosphatase zeta (PTPRZ_HUMAN) | P23471 | | 7 | PTPRZ1 | Homo sapiens | 9606 | 1.3 | 1.3 | 54.95 | 1 | 1 | 1 | 1 | 2 | 1 | | | | | 22.30752849 | 0.000635114 | 254.4290053 | 100 | Doubtful |
| 12 | Single Protein | P01023 | Alpha-2-macroglobulin (A2MG_HUMAN) | P01023 | | 12 | A2M | Homo sapiens | 9606 | 5.29 | 5.29 | 70.96 | 3 | 3 | 3 | 3 | 8 | 5 | | | | | 135.6365977 | 0.00386169 | 163.1878915 | 100 | Doubtful |
| | | | | | | | | | | | | | | | | | | | | | | total fibrinogen | 11617.21012 | 0.330751881 | | | |
