## Supplementary for "High-Throughput Site-Specific N-Glycosylation Profiling of Human Fibrinogen in Atrial Fibrillation": Table S3.pdf

**Table S3.** Intra-plate (8 replicates were randomized across 1 plate) and inter-plate (16 replicates were randomized across 4 different plates) repeatability of the reported glycopeptides. StDev – standard deviation

| Standard | Plate | Beta.normal-N4H5S1 | Beta.normal-N4H5S2 | Beta.MISS-N4H5S1 | Beta.MISS-N4H5S2 | Gamma.normal-N4H5S1 | Gamma.normal-N4H5S2 | Gamma.MISS-N4H5 | Gamma.MISS-N4H5S1 | Gamma.MISS-N4H5S1F1 | Gamma.MISS-N4H5S2 | Gamma.MISS-N4H5S2F1 | Average | StDev |
|---|---|---|---|---|---|---|---|---|---|---|---|---|---|---|
| stand_01 | 1 | 55.7 | 44.3 | 55.2 | 44.8 | 55.5 | 44.5 | 1.5 | 69.1 | 1.9 | 26.6 | 0.8 | | |
| stand_02 | 1 | 55.7 | 44.3 | 57.4 | 42.6 | 54.6 | 45.4 | 1.5 | 69.2 | 1.9 | 26.7 | 0.8 | | |
| stand_03 | 1 | 55.3 | 44.7 | 56.8 | 43.2 | 54.1 | 45.9 | 1.5 | 68.8 | 1.9 | 27.0 | 0.8 | | |
| stand_04 | 1 | 54.4 | 45.6 | 59.6 | 40.4 | 49.8 | 50.2 | 1.4 | 69.2 | 1.8 | 26.9 | 0.7 | | |
| stand_05 | 1 | 55.2 | 44.8 | 59.5 | 40.5 | 54.8 | 45.2 | 1.5 | 68.7 | 1.8 | 27.3 | 0.7 | | |
| stand_06 | 1 | 55.3 | 44.7 | 57.8 | 42.2 | 54.0 | 46.0 | 1.5 | 68.8 | 1.9 | 27.0 | 0.8 | | |
| stand_07 | 1 | 55.2 | 44.8 | 59.3 | 40.7 | 52.6 | 47.4 | 1.6 | 68.7 | 1.8 | 27.1 | 0.7 | | |
| stand_08 | 1 | 54.0 | 46.0 | 59.3 | 40.7 | 41.1 | 58.9 | 2.0 | 67.0 | 1.7 | 28.5 | 0.7 | | |
| stand_09 | 2 | 54.1 | 45.9 | 59.0 | 41.0 | 53.2 | 46.8 | 2.2 | 68.3 | 1.8 | 27.0 | 0.7 | | |
| stand_10 | 2 | 50.5 | 49.5 | 57.8 | 42.2 | 34.5 | 65.5 | 3.6 | 67.5 | 1.5 | 26.5 | 0.9 | | |
| stand_11 | 3 | 57.2 | 42.8 | 58.4 | 41.6 | 55.4 | 44.6 | 2.5 | 67.8 | 1.7 | 27.3 | 0.7 | | |
| stand_12 | 3 | 58.8 | 41.2 | 59.1 | 40.9 | 53.3 | 46.7 | 2.1 | 67.9 | 1.6 | 27.8 | 0.6 | | |
| stand_13 | 3 | 59.8 | 40.2 | 59.0 | 41.0 | 53.8 | 46.2 | 1.9 | 68.4 | 1.5 | 27.6 | 0.6 | | |
| stand_14 | 4 | 62.9 | 37.1 | 63.0 | 37.0 | 55.4 | 44.6 | 2.0 | 70.2 | 1.5 | 25.6 | 0.7 | | |
| stand_15 | 4 | 56.7 | 43.3 | 63.9 | 36.1 | 54.0 | 46.0 | 1.8 | 69.1 | 1.8 | 26.5 | 0.8 | | |
| stand_16 | 4 | 55.6 | 44.4 | 60.0 | 40.0 | 30.8 | 69.2 | 3.0 | 66.8 | 2.5 | 25.7 | 1.9 | | |
| **CV (intra-plate)** | | 1.1% | 1.3% | 2.8% | 3.8% | 9.2% | 10.0% | 12.5% | 1.0% | 4.7% | 2.2% | 5.4% | **4.9%** | 4.0% |
| **CV (inter-plate)** | | 4.9% | 6.3% | 3.6% | 5.2% | 15.5% | 15.7% | 31.4% | 1.3% | 13.4% | 2.6% | 37.0% | **12.4%** | 12.0% |
