## Supplementary for "High-Throughput Site-Specific N-Glycosylation Profiling of Human Fibrinogen in Atrial Fibrillation": Table S4.pdf

**Table S4.** Assocciation of fibrinogen glycopeptides with age and sex. Std.error – standard error, p.adj_LJ – false discovery rate was controlled using Benjamini–Hochberg method modified by Li and Ji at the specified level of 0.05.

| Glycopeptide | Variable | Estimate | Std.error | p.value | p.adj_LJ |
|---|---|---|---|---|---|
| Beta.normal-N4H5S2 | Age | -0.0307 | 0.0075 | 0.0001 | 0.0003 |
| Beta.normal-N4H5S1 | Age | 0.0288 | 0.0075 | 0.0002 | 0.0005 |
| Gamma.MISS-N4H5S2 | Age | -0.0212 | 0.0072 | 0.0036 | 0.0071 |
| Gamma.MISS-N4H5S1 | Age | 0.0134 | 0.0072 | 0.0640 | 0.1025 |
| Gamma.MISS-N4H5 | Age | 0.0087 | 0.0066 | 0.1930 | 0.2383 |
| Gamma.MISS-N4H5S1F1 | Age | 0.0086 | 0.0068 | 0.2085 | 0.2383 |
| Gamma.MISS-N4H5S2F1 | Age | 0.0009 | 0.0067 | 0.8903 | 0.8903 |
| Beta.normal-N4H5S2 | SexM | 0.4095 | 0.1600 | 0.0114 | 0.0307 |
| Gamma.MISS-N4H5S1F1 | SexM | -0.3101 | 0.1268 | 0.0153 | 0.0307 |
| Gamma.MISS-N4H5S2 | SexM | 0.3424 | 0.1365 | 0.0129 | 0.0307 |
| Beta.normal-N4H5S1 | SexM | -0.3539 | 0.1613 | 0.0297 | 0.0475 |
| Gamma.MISS-N4H5 | SexM | -0.1955 | 0.1245 | 0.1181 | 0.1444 |
| Gamma.MISS-N4H5S2F1 | SexM | -0.1907 | 0.1242 | 0.1263 | 0.1444 |
| Gamma.MISS-N4H5S1 | SexM | -0.1183 | 0.1366 | 0.3872 | 0.3872 |
