## Supplementary for "High-Throughput Site-Specific N-Glycosylation Profiling of Human Fibrinogen in Atrial Fibrillation": Table S5.pdf

**Table S5.** Potential differences in fibrinogen glycosylation between patients who experienced AF recurrence within six months post-catheter ablation and those who did not. Initial – glycan profiles of patients with AF recurrence versus those without at the initial time point, Follow up – profiles of recurrence versus non-recurrence at the six-month follow-up, Delta – changes in glycan profiles over time between the two time points for patients with and without recurrence, Std.error – standard error, p.adj_LJ – false discovery rate was controlled using Benjamini–Hochberg method modified by Li and Ji at the specified level of 0.05.

| Glycopeptide | Timepoint | Estimate | Std.error | p.value | p.adj_LJ |
|---|---|---|---|---|---|
| Beta.normal-N4H5S1 | initial | 0.125 | 0.227 | 0.583 | 0.999 |
| Beta.normal-N4H5S1 | follow-up | 0.094 | 0.262 | 0.722 | 0.999 |
| Beta.normal-N4H5S2 | initial | -0.130 | 0.227 | 0.569 | 0.999 |
| Beta.normal-N4H5S2 | follow-up | -0.100 | 0.261 | 0.702 | 0.999 |
| Gamma.MISS-N4H5 | initial | -0.098 | 0.187 | 0.601 | 0.999 |
| Gamma.MISS-N4H5 | follow-up | -0.068 | 0.222 | 0.759 | 0.999 |
| Gamma.MISS-N4H5S1 | initial | 0.148 | 0.184 | 0.423 | 0.999 |
| Gamma.MISS-N4H5S1 | follow-up | 0.000 | 0.209 | 0.999 | 0.999 |
| Gamma.MISS-N4H5S1F1 | initial | 0.026 | 0.186 | 0.891 | 0.999 |
| Gamma.MISS-N4H5S1F1 | follow-up | 0.075 | 0.221 | 0.735 | 0.999 |
| Gamma.MISS-N4H5S2 | initial | -0.010 | 0.179 | 0.955 | 0.999 |
| Gamma.MISS-N4H5S2 | follow-up | 0.010 | 0.194 | 0.960 | 0.999 |
| Gamma.MISS-N4H5S2F1 | initial | -0.017 | 0.187 | 0.926 | 0.999 |
| Gamma.MISS-N4H5S2F1 | follow-up | 0.199 | 0.223 | 0.372 | 0.999 |
| Beta.normal-N4H5S1 | delta | -0.032 | 0.262 | 0.904 | 0.912 |
| Beta.normal-N4H5S2 | delta | 0.030 | 0.260 | 0.909 | 0.912 |
| Gamma.MISS-N4H5 | delta | 0.030 | 0.268 | 0.912 | 0.912 |
| Gamma.MISS-N4H5S1 | delta | -0.147 | 0.209 | 0.483 | 0.912 |
| Gamma.MISS-N4H5S1F1 | delta | 0.049 | 0.260 | 0.851 | 0.912 |
| Gamma.MISS-N4H5S2 | delta | 0.020 | 0.149 | 0.895 | 0.912 |
| Gamma.MISS-N4H5S2F1 | delta | 0.217 | 0.279 | 0.438 | 0.912 |
