## Supplementary for "High-Throughput Site-Specific N-Glycosylation Profiling of Human Fibrinogen in Atrial Fibrillation": Table S6.pdf

**Table S6.** Assocciation of fibrinogen glycopeptides with biochemical, hematological parameters, as well as BMI. Std.error – standard error, p.adj_LJ – false discovery rate was controlled using Benjamini–Hochberg method modified by Li and Ji at the specified level of 0.05.

| Variable | Group | Glycopeptide | Estimate | Std.error | p.value | p.adj_LJ |
|---|---|---|---|---|---|---|
| Basophils (relative) | Blood Cell Counts | Beta.normal-N4H5S2 | 0.27469 | 0.14871 | 0.07296 | 0.18209 |
| Basophils (relative) | Blood Cell Counts | Gamma.MISS-N4H5S2F1 | 0.24585 | 0.14229 | 0.09104 | 0.18209 |
| Basophils (relative) | Blood Cell Counts | Beta.normal-N4H5S1 | -0.29047 | 0.15316 | 0.06593 | 0.18209 |
| Basophils (relative) | Blood Cell Counts | Gamma.MISS-N4H5S1F1 | 0.20592 | 0.14227 | 0.15488 | 0.24781 |
| Basophils (relative) | Blood Cell Counts | Gamma.MISS-N4H5S1 | -0.18207 | 0.14959 | 0.23003 | 0.30670 |
| Basophils (relative) | Blood Cell Counts | Gamma.MISS-N4H5S2 | 0.10982 | 0.14844 | 0.46334 | 0.52953 |
| Basophils (relative) | Blood Cell Counts | Gamma.MISS-N4H5 | 0.03853 | 0.14170 | 0.78697 | 0.78697 |
| Body Mass Index | Biochemical Parameters and BMI | Gamma.MISS-N4H5S2 | 0.32579 | 0.07774 | 0.00005 | 0.00019 |
| Body Mass Index | Biochemical Parameters and BMI | Beta.normal-N4H5S1 | -0.29966 | 0.09799 | 0.00290 | 0.00581 |
| Body Mass Index | Biochemical Parameters and BMI | Beta.normal-N4H5S2 | 0.30137 | 0.09767 | 0.00267 | 0.00581 |
| Body Mass Index | Biochemical Parameters and BMI | Gamma.MISS-N4H5S1 | -0.19198 | 0.08215 | 0.02085 | 0.03335 |
| Body Mass Index | Biochemical Parameters and BMI | Gamma.MISS-N4H5 | -0.10061 | 0.08339 | 0.22959 | 0.30612 |
| Body Mass Index | Biochemical Parameters and BMI | Gamma.MISS-N4H5S2F1 | 0.06043 | 0.08384 | 0.47227 | 0.53973 |
| Body Mass Index | Biochemical Parameters and BMI | Gamma.MISS-N4H5S1F1 | 0.02063 | 0.08337 | 0.80495 | 0.80495 |
| B-Type Natriuretic Peptide | Biochemical Parameters and BMI | Gamma.MISS-N4H5 | -0.15339 | 0.09589 | 0.11204 | 0.44816 |
| B-Type Natriuretic Peptide | Biochemical Parameters and BMI | Gamma.MISS-N4H5S2F1 | -0.11012 | 0.09659 | 0.25630 | 0.68347 |
| B-Type Natriuretic Peptide | Biochemical Parameters and BMI | Beta.normal-N4H5S2 | -0.08859 | 0.11899 | 0.45858 | 0.73373 |
| B-Type Natriuretic Peptide | Biochemical Parameters and BMI | Beta.normal-N4H5S1 | 0.10604 | 0.11918 | 0.37608 | 0.73373 |
| B-Type Natriuretic Peptide | Biochemical Parameters and BMI | Gamma.MISS-N4H5S1 | 0.04980 | 0.09669 | 0.60738 | 0.80984 |
| B-Type Natriuretic Peptide | Biochemical Parameters and BMI | Gamma.MISS-N4H5S1F1 | 0.02801 | 0.09676 | 0.77264 | 0.88302 |
| B-Type Natriuretic Peptide | Biochemical Parameters and BMI | Gamma.MISS-N4H5S2 | -0.00602 | 0.09554 | 0.94986 | 0.94986 |
| Cholesterol | Biochemical Parameters and BMI | Gamma.MISS-N4H5S1F1 | -0.08450 | 0.07893 | 0.28600 | 0.68613 |
| Cholesterol | Biochemical Parameters and BMI | Beta.normal-N4H5S2 | 0.08662 | 0.09097 | 0.34307 | 0.68613 |
| Cholesterol | Biochemical Parameters and BMI | Beta.normal-N4H5S1 | -0.09305 | 0.09093 | 0.30837 | 0.68613 |
| Cholesterol | Biochemical Parameters and BMI | Gamma.MISS-N4H5S1 | -0.03538 | 0.08019 | 0.65969 | 0.78266 |
| Cholesterol | Biochemical Parameters and BMI | Gamma.MISS-N4H5S2 | 0.02960 | 0.07956 | 0.71035 | 0.78266 |
| Cholesterol | Biochemical Parameters and BMI | Gamma.MISS-N4H5S2F1 | -0.05273 | 0.07886 | 0.50466 | 0.78266 |
| Cholesterol | Biochemical Parameters and BMI | Gamma.MISS-N4H5 | -0.02151 | 0.07786 | 0.78266 | 0.78266 |
| Creatinine | Biochemical Parameters and BMI | Gamma.MISS-N4H5S1F1 | 0.18580 | 0.08650 | 0.03320 | 0.13279 |
| Creatinine | Biochemical Parameters and BMI | Gamma.MISS-N4H5S2F1 | 0.08763 | 0.08771 | 0.31922 | 0.69773 |
| Creatinine | Biochemical Parameters and BMI | Beta.normal-N4H5S2 | 0.09345 | 0.09935 | 0.34887 | 0.69773 |
| Creatinine | Biochemical Parameters and BMI | Gamma.MISS-N4H5S2 | 0.04694 | 0.08672 | 0.58913 | 0.69997 |
| Creatinine | Biochemical Parameters and BMI | Beta.normal-N4H5S1 | -0.06831 | 0.10057 | 0.49838 | 0.69997 |
| Creatinine | Biochemical Parameters and BMI | Gamma.MISS-N4H5S1 | -0.04414 | 0.08697 | 0.61247 | 0.69997 |
| Creatinine | Biochemical Parameters and BMI | Gamma.MISS-N4H5 | -0.02711 | 0.08629 | 0.75374 | 0.75374 |
| C-Reactive Protein | Biochemical Parameters and BMI | Gamma.MISS-N4H5S1F1 | -0.04919 | 0.14318 | 0.73180 | 0.98534 |
| C-Reactive Protein | Biochemical Parameters and BMI | Gamma.MISS-N4H5 | -0.02285 | 0.14393 | 0.87415 | 0.98534 |
| C-Reactive Protein | Biochemical Parameters and BMI | Gamma.MISS-N4H5S2F1 | -0.03891 | 0.14363 | 0.78695 | 0.98534 |
| C-Reactive Protein | Biochemical Parameters and BMI | Gamma.MISS-N4H5S1 | 0.06180 | 0.14237 | 0.66505 | 0.98534 |
| C-Reactive Protein | Biochemical Parameters and BMI | Beta.normal-N4H5S1 | 0.05834 | 0.17591 | 0.74110 | 0.98534 |
| C-Reactive Protein | Biochemical Parameters and BMI | Beta.normal-N4H5S2 | -0.05121 | 0.17539 | 0.77110 | 0.98534 |
| C-Reactive Protein | Biochemical Parameters and BMI | Gamma.MISS-N4H5S2 | 0.00260 | 0.14145 | 0.98534 | 0.98534 |
| Eosinophils (relative) | Blood Cell Counts | Gamma.MISS-N4H5S2F1 | 0.23526 | 0.14440 | 0.11040 | 0.27002 |
| Eosinophils (relative) | Blood Cell Counts | Beta.normal-N4H5S2 | 0.23572 | 0.15417 | 0.13501 | 0.27002 |
| Eosinophils (relative) | Blood Cell Counts | Gamma.MISS-N4H5S1F1 | 0.21908 | 0.14350 | 0.13400 | 0.27002 |
| Eosinophils (relative) | Blood Cell Counts | Beta.normal-N4H5S1 | -0.17842 | 0.16152 | 0.27665 | 0.44263 |
| Eosinophils (relative) | Blood Cell Counts | Gamma.MISS-N4H5S1 | -0.06535 | 0.15347 | 0.67232 | 0.88628 |
| Eosinophils (relative) | Blood Cell Counts | Gamma.MISS-N4H5 | -0.02278 | 0.14337 | 0.87447 | 0.88628 |
| Eosinophils (relative) | Blood Cell Counts | Gamma.MISS-N4H5S2 | 0.02172 | 0.15100 | 0.88628 | 0.88628 |
| Erythroctytes | Blood Cell Counts | Gamma.MISS-N4H5 | -0.34655 | 0.17419 | 0.05288 | 0.21153 |
| Erythroctytes | Blood Cell Counts | Gamma.MISS-N4H5S1F1 | -0.24165 | 0.18314 | 0.19383 | 0.51688 |
| Erythroctytes | Blood Cell Counts | Gamma.MISS-N4H5S2 | 0.16000 | 0.19000 | 0.40427 | 0.80855 |
| Erythroctytes | Blood Cell Counts | Gamma.MISS-N4H5S1 | -0.10753 | 0.19434 | 0.58286 | 0.82161 |
| Erythroctytes | Blood Cell Counts | Beta.normal-N4H5S1 | -0.10924 | 0.21605 | 0.61620 | 0.82161 |
| Erythroctytes | Blood Cell Counts | Beta.normal-N4H5S2 | 0.06273 | 0.20977 | 0.76664 | 0.87616 |
| Erythroctytes | Blood Cell Counts | Gamma.MISS-N4H5S2F1 | -0.00742 | 0.18855 | 0.96878 | 0.96878 |
| Glucose | Biochemical Parameters and BMI | Gamma.MISS-N4H5S2 | 0.22168 | 0.07596 | 0.00401 | 0.01603 |
| Glucose | Biochemical Parameters and BMI | Gamma.MISS-N4H5S1 | -0.16267 | 0.07769 | 0.03781 | 0.10083 |
| Glucose | Biochemical Parameters and BMI | Gamma.MISS-N4H5S2F1 | 0.14302 | 0.07905 | 0.07222 | 0.13769 |
| Glucose | Biochemical Parameters and BMI | Beta.normal-N4H5S2 | 0.14827 | 0.09028 | 0.10327 | 0.13769 |
| Glucose | Biochemical Parameters and BMI | Gamma.MISS-N4H5S1F1 | 0.13594 | 0.07893 | 0.08688 | 0.13769 |
| Glucose | Biochemical Parameters and BMI | Beta.normal-N4H5S1 | -0.11974 | 0.09165 | 0.19401 | 0.22173 |
| Glucose | Biochemical Parameters and BMI | Gamma.MISS-N4H5 | -0.08129 | 0.07788 | 0.29816 | 0.29816 |
| Hemoglobin | Blood Cell Counts | Gamma.MISS-N4H5 | -0.14963 | 0.09448 | 0.11517 | 0.21007 |
| Hemoglobin | Blood Cell Counts | Beta.normal-N4H5S1 | -0.18302 | 0.11856 | 0.12537 | 0.21007 |
| Hemoglobin | Blood Cell Counts | Beta.normal-N4H5S2 | 0.17717 | 0.11738 | 0.13393 | 0.21007 |
| Hemoglobin | Blood Cell Counts | Gamma.MISS-N4H5S1F1 | -0.13674 | 0.09631 | 0.15755 | 0.21007 |
| Hemoglobin | Blood Cell Counts | Gamma.MISS-N4H5S2 | 0.17817 | 0.09519 | 0.06302 | 0.21007 |
| Hemoglobin | Blood Cell Counts | Gamma.MISS-N4H5S1 | -0.08117 | 0.09699 | 0.40385 | 0.46154 |
| Hemoglobin | Blood Cell Counts | Gamma.MISS-N4H5S2F1 | -0.03198 | 0.09704 | 0.74213 | 0.74213 |
| HDL Cholesterol | Biochemical Parameters and BMI | Beta.normal-N4H5S2 | -0.38843 | 0.15076 | 0.01436 | 0.04808 |
| HDL Cholesterol | Biochemical Parameters and BMI | Beta.normal-N4H5S1 | 0.37641 | 0.15168 | 0.01803 | 0.04808 |
| HDL Cholesterol | Biochemical Parameters and BMI | Gamma.MISS-N4H5S2 | -0.29528 | 0.14249 | 0.04427 | 0.08854 |
| HDL Cholesterol | Biochemical Parameters and BMI | Gamma.MISS-N4H5S1 | 0.27196 | 0.14765 | 0.07239 | 0.11582 |
| HDL Cholesterol | Biochemical Parameters and BMI | Gamma.MISS-N4H5S1F1 | -0.18691 | 0.14494 | 0.20408 | 0.27211 |
| HDL Cholesterol | Biochemical Parameters and BMI | Gamma.MISS-N4H5S2F1 | -0.15859 | 0.14743 | 0.28806 | 0.32921 |
| HDL Cholesterol | Biochemical Parameters and BMI | Gamma.MISS-N4H5 | 0.13983 | 0.14205 | 0.33043 | 0.33043 |
| Hematocrit | Erythrocyte Indices | Gamma.MISS-N4H5 | -0.45867 | 0.16314 | 0.00734 | 0.02935 |
| Hematocrit | Erythrocyte Indices | Gamma.MISS-N4H5S2 | 0.27223 | 0.18205 | 0.14195 | 0.37853 |
| Hematocrit | Erythrocyte Indices | Beta.normal-N4H5S1 | -0.21665 | 0.21281 | 0.31545 | 0.57711 |
| Hematocrit | Erythrocyte Indices | Gamma.MISS-N4H5S1 | -0.17384 | 0.18820 | 0.36069 | 0.57711 |
| Hematocrit | Erythrocyte Indices | Beta.normal-N4H5S2 | 0.12014 | 0.20814 | 0.56741 | 0.65290 |
| Hematocrit | Erythrocyte Indices | Gamma.MISS-N4H5S1F1 | -0.10341 | 0.18127 | 0.57128 | 0.65290 |
| Hematocrit | Erythrocyte Indices | Gamma.MISS-N4H5S2F1 | 0.06052 | 0.18349 | 0.74309 | 0.74309 |
| Potassium | Biochemical Parameters and BMI | Gamma.MISS-N4H5S1F1 | 0.15601 | 0.09168 | 0.09147 | 0.36586 |
| Potassium | Biochemical Parameters and BMI | Gamma.MISS-N4H5S1 | -0.04263 | 0.09204 | 0.64410 | 0.82053 |
| Potassium | Biochemical Parameters and BMI | Beta.normal-N4H5S2 | 0.04041 | 0.11147 | 0.71797 | 0.82053 |
| Potassium | Biochemical Parameters and BMI | Gamma.MISS-N4H5 | -0.08072 | 0.09284 | 0.38640 | 0.82053 |
| Potassium | Biochemical Parameters and BMI | Gamma.MISS-N4H5S2F1 | 0.04135 | 0.09299 | 0.65737 | 0.82053 |
| Potassium | Biochemical Parameters and BMI | Beta.normal-N4H5S1 | -0.04989 | 0.11179 | 0.65667 | 0.82053 |
| Potassium | Biochemical Parameters and BMI | Gamma.MISS-N4H5S2 | 0.01388 | 0.09136 | 0.87949 | 0.87949 |
| LDL Cholesterol | Biochemical Parameters and BMI | Gamma.MISS-N4H5S1F1 | -0.10395 | 0.07850 | 0.18726 | 0.37451 |
| LDL Cholesterol | Biochemical Parameters and BMI | Beta.normal-N4H5S1 | -0.12528 | 0.08950 | 0.16432 | 0.37451 |
| LDL Cholesterol | Biochemical Parameters and BMI | Beta.normal-N4H5S2 | 0.12152 | 0.08935 | 0.17650 | 0.37451 |
| LDL Cholesterol | Biochemical Parameters and BMI | Gamma.MISS-N4H5S1 | -0.06273 | 0.07900 | 0.42832 | 0.68531 |
| LDL Cholesterol | Biochemical Parameters and BMI | Gamma.MISS-N4H5S2 | 0.04820 | 0.07818 | 0.53841 | 0.71788 |
| LDL Cholesterol | Biochemical Parameters and BMI | Gamma.MISS-N4H5S2F1 | -0.02950 | 0.07918 | 0.70997 | 0.81140 |
| LDL Cholesterol | Biochemical Parameters and BMI | Gamma.MISS-N4H5 | 0.01841 | 0.07780 | 0.81320 | 0.81320 |
| Leukocytes | Blood Cell Counts | Gamma.MISS-N4H5S2 | -0.10267 | 0.14431 | 0.48053 | 0.92163 |
| Leukocytes | Blood Cell Counts | Gamma.MISS-N4H5S1F1 | 0.07944 | 0.14099 | 0.57602 | 0.92163 |
| Leukocytes | Blood Cell Counts | Gamma.MISS-N4H5S1 | 0.11321 | 0.14679 | 0.44469 | 0.92163 |
| Leukocytes | Blood Cell Counts | Beta.normal-N4H5S2 | -0.09310 | 0.15241 | 0.54515 | 0.92163 |
| Leukocytes | Blood Cell Counts | Beta.normal-N4H5S1 | 0.04562 | 0.15796 | 0.77440 | 0.98721 |
| Leukocytes | Blood Cell Counts | Gamma.MISS-N4H5S2F1 | 0.00230 | 0.14288 | 0.98721 | 0.98721 |
| Leukocytes | Blood Cell Counts | Gamma.MISS-N4H5 | 0.02058 | 0.13777 | 0.88193 | 0.98721 |
| Lymphocytes (relative) | Blood Cell Counts | Gamma.MISS-N4H5S2 | 0.11987 | 0.14923 | 0.42612 | 0.71153 |
| Lymphocytes (relative) | Blood Cell Counts | Gamma.MISS-N4H5 | -0.07483 | 0.14228 | 0.60157 | 0.71153 |
| Lymphocytes (relative) | Blood Cell Counts | Gamma.MISS-N4H5S1F1 | 0.11721 | 0.14549 | 0.42479 | 0.71153 |
| Lymphocytes (relative) | Blood Cell Counts | Gamma.MISS-N4H5S1 | -0.09724 | 0.15236 | 0.52664 | 0.71153 |
| Lymphocytes (relative) | Blood Cell Counts | Beta.normal-N4H5S1 | -0.11045 | 0.15768 | 0.48813 | 0.71153 |
| Lymphocytes (relative) | Blood Cell Counts | Beta.normal-N4H5S2 | 0.07609 | 0.15326 | 0.62258 | 0.71153 |
| Lymphocytes (relative) | Blood Cell Counts | Gamma.MISS-N4H5S2F1 | 0.02871 | 0.14792 | 0.84700 | 0.84700 |
| Large Unstained Cells (relative) | Blood Cell Counts | Beta.normal-N4H5S1 | -0.17926 | 0.16646 | 0.28890 | 0.50218 |
| Large Unstained Cells (relative) | Blood Cell Counts | Gamma.MISS-N4H5 | -0.14138 | 0.13973 | 0.31731 | 0.50218 |
| Large Unstained Cells (relative) | Blood Cell Counts | Gamma.MISS-N4H5S2 | 0.20156 | 0.14697 | 0.17735 | 0.50218 |
| Large Unstained Cells (relative) | Blood Cell Counts | Gamma.MISS-N4H5S1 | -0.19070 | 0.15006 | 0.21063 | 0.50218 |
| Large Unstained Cells (relative) | Blood Cell Counts | Gamma.MISS-N4H5S1F1 | 0.11333 | 0.14522 | 0.43941 | 0.50218 |
| Large Unstained Cells (relative) | Blood Cell Counts | Beta.normal-N4H5S2 | 0.12924 | 0.16245 | 0.43164 | 0.50218 |
| Large Unstained Cells (relative) | Blood Cell Counts | Gamma.MISS-N4H5S2F1 | 0.03267 | 0.14735 | 0.82559 | 0.82559 |
| Mean Corpuscular Hemoglobin | Erythrocyte Indices | Gamma.MISS-N4H5S1F1 | 0.34632 | 0.13606 | 0.01450 | 0.05799 |
| Mean Corpuscular Hemoglobin | Erythrocyte Indices | Gamma.MISS-N4H5S2F1 | 0.18584 | 0.14446 | 0.20501 | 0.30044 |
| Mean Corpuscular Hemoglobin | Erythrocyte Indices | Gamma.MISS-N4H5S2 | 0.22297 | 0.14565 | 0.13294 | 0.30044 |
| Mean Corpuscular Hemoglobin | Erythrocyte Indices | Beta.normal-N4H5S2 | 0.19893 | 0.16125 | 0.22533 | 0.30044 |
| Mean Corpuscular Hemoglobin | Erythrocyte Indices | Gamma.MISS-N4H5S1 | -0.21483 | 0.14872 | 0.15567 | 0.30044 |
| Mean Corpuscular Hemoglobin | Erythrocyte Indices | Beta.normal-N4H5S1 | -0.18231 | 0.16720 | 0.28279 | 0.32319 |
| Mean Corpuscular Hemoglobin | Erythrocyte Indices | Gamma.MISS-N4H5 | -0.11190 | 0.14092 | 0.43143 | 0.43143 |
| Mean Corpuscular Hemoglobin Concentration | Erythrocyte Indices | Beta.normal-N4H5S2 | 0.38848 | 0.17438 | 0.03223 | 0.09907 |
| Mean Corpuscular Hemoglobin Concentration | Erythrocyte Indices | Gamma.MISS-N4H5S1F1 | 0.32849 | 0.15981 | 0.04579 | 0.09907 |
| Mean Corpuscular Hemoglobin Concentration | Erythrocyte Indices | Gamma.MISS-N4H5S1 | -0.33760 | 0.16716 | 0.04954 | 0.09907 |
| Mean Corpuscular Hemoglobin Concentration | Erythrocyte Indices | Gamma.MISS-N4H5S2 | 0.28522 | 0.16613 | 0.09304 | 0.12918 |
| Mean Corpuscular Hemoglobin Concentration | Erythrocyte Indices | Beta.normal-N4H5S1 | -0.31485 | 0.18471 | 0.09689 | 0.12918 |
| Mean Corpuscular Hemoglobin Concentration | Erythrocyte Indices | Gamma.MISS-N4H5S2F1 | 0.25652 | 0.16445 | 0.12596 | 0.14396 |
| Mean Corpuscular Hemoglobin Concentration | Erythrocyte Indices | Gamma.MISS-N4H5 | -0.05948 | 0.16269 | 0.71639 | 0.71639 |
| Mean Corpuscular Volume | Erythrocyte Indices | Gamma.MISS-N4H5S1F1 | 0.23361 | 0.13914 | 0.10024 | 0.40097 |
| Mean Corpuscular Volume | Erythrocyte Indices | Gamma.MISS-N4H5S2F1 | 0.09038 | 0.14428 | 0.53430 | 0.88768 |
| Mean Corpuscular Volume | Erythrocyte Indices | Beta.normal-N4H5S1 | -0.02298 | 0.16153 | 0.88768 | 0.88768 |
| Mean Corpuscular Volume | Erythrocyte Indices | Gamma.MISS-N4H5 | -0.09985 | 0.13896 | 0.47621 | 0.88768 |
| Mean Corpuscular Volume | Erythrocyte Indices | Gamma.MISS-N4H5S1 | -0.04115 | 0.14977 | 0.78479 | 0.88768 |
| Mean Corpuscular Volume | Erythrocyte Indices | Gamma.MISS-N4H5S2 | 0.07654 | 0.14676 | 0.60460 | 0.88768 |
| Mean Corpuscular Volume | Erythrocyte Indices | Beta.normal-N4H5S2 | -0.03780 | 0.15639 | 0.81041 | 0.88768 |
| Monocytes (relative) | Blood Cell Counts | Beta.normal-N4H5S1 | -0.23651 | 0.16313 | 0.15577 | 0.62307 |
| Monocytes (relative) | Blood Cell Counts | Beta.normal-N4H5S2 | 0.18319 | 0.15968 | 0.25886 | 0.69029 |
| Monocytes (relative) | Blood Cell Counts | Gamma.MISS-N4H5S1F1 | 0.12275 | 0.14596 | 0.40490 | 0.80980 |
| Monocytes (relative) | Blood Cell Counts | Gamma.MISS-N4H5S2 | 0.02064 | 0.15087 | 0.89179 | 0.96874 |
| Monocytes (relative) | Blood Cell Counts | Gamma.MISS-N4H5 | -0.04334 | 0.14313 | 0.76350 | 0.96874 |
| Monocytes (relative) | Blood Cell Counts | Gamma.MISS-N4H5S1 | -0.00606 | 0.15365 | 0.96874 | 0.96874 |
| Monocytes (relative) | Blood Cell Counts | Gamma.MISS-N4H5S2F1 | -0.01880 | 0.14853 | 0.89987 | 0.96874 |
| Mean Platelet Volume | Erythrocyte Indices | Gamma.MISS-N4H5 | -0.23736 | 0.13328 | 0.08183 | 0.32734 |
| Mean Platelet Volume | Erythrocyte Indices | Gamma.MISS-N4H5S2F1 | -0.18426 | 0.14036 | 0.19606 | 0.52283 |
| Mean Platelet Volume | Erythrocyte Indices | Gamma.MISS-N4H5S2 | -0.05847 | 0.14507 | 0.68888 | 0.85991 |
| Mean Platelet Volume | Erythrocyte Indices | Gamma.MISS-N4H5S1F1 | -0.04757 | 0.14152 | 0.73838 | 0.85991 |
| Mean Platelet Volume | Erythrocyte Indices | Beta.normal-N4H5S2 | 0.04913 | 0.15457 | 0.75242 | 0.85991 |
| Mean Platelet Volume | Erythrocyte Indices | Gamma.MISS-N4H5S1 | 0.09735 | 0.14726 | 0.51201 | 0.85991 |
| Mean Platelet Volume | Erythrocyte Indices | Beta.normal-N4H5S1 | 0.00837 | 0.15978 | 0.95851 | 0.95851 |
| Neutrophils (relative) | Blood Cell Counts | Beta.normal-N4H5S1 | 0.11875 | 0.15799 | 0.45717 | 0.86091 |
| Neutrophils (relative) | Blood Cell Counts | Gamma.MISS-N4H5S1F1 | -0.13536 | 0.14248 | 0.34731 | 0.86091 |
| Neutrophils (relative) | Blood Cell Counts | Beta.normal-N4H5S2 | -0.08240 | 0.15364 | 0.59505 | 0.86091 |
| Neutrophils (relative) | Blood Cell Counts | Gamma.MISS-N4H5 | 0.06474 | 0.13984 | 0.64568 | 0.86091 |
| Neutrophils (relative) | Blood Cell Counts | Gamma.MISS-N4H5S2 | -0.06852 | 0.14727 | 0.64404 | 0.86091 |
| Neutrophils (relative) | Blood Cell Counts | Gamma.MISS-N4H5S1 | 0.04270 | 0.15019 | 0.77750 | 0.88857 |
| Neutrophils (relative) | Blood Cell Counts | Gamma.MISS-N4H5S2F1 | -0.01498 | 0.14532 | 0.91837 | 0.91837 |
| Red Cell Distribution Width | Erythrocyte Indices | Gamma.MISS-N4H5S1 | 0.12994 | 0.16664 | 0.43969 | 0.89753 |
| Red Cell Distribution Width | Erythrocyte Indices | Gamma.MISS-N4H5 | -0.02026 | 0.15643 | 0.89753 | 0.89753 |
| Red Cell Distribution Width | Erythrocyte Indices | Beta.normal-N4H5S1 | 0.11833 | 0.19235 | 0.54231 | 0.89753 |
| Red Cell Distribution Width | Erythrocyte Indices | Gamma.MISS-N4H5S2F1 | -0.05861 | 0.16198 | 0.71921 | 0.89753 |
| Red Cell Distribution Width | Erythrocyte Indices | Gamma.MISS-N4H5S2 | -0.08142 | 0.16432 | 0.62274 | 0.89753 |
| Red Cell Distribution Width | Erythrocyte Indices | Gamma.MISS-N4H5S1F1 | 0.03737 | 0.16056 | 0.81702 | 0.89753 |
| Red Cell Distribution Width | Erythrocyte Indices | Beta.normal-N4H5S2 | -0.12068 | 0.18623 | 0.52110 | 0.89753 |
| Thrombocytes | Blood Cell Counts | Gamma.MISS-N4H5 | 0.13919 | 0.13930 | 0.32316 | 0.94815 |
| Thrombocytes | Blood Cell Counts | Beta.normal-N4H5S2 | 0.01054 | 0.16097 | 0.94815 | 0.94815 |
| Thrombocytes | Blood Cell Counts | Gamma.MISS-N4H5S2F1 | 0.08049 | 0.14555 | 0.58305 | 0.94815 |
| Thrombocytes | Blood Cell Counts | Gamma.MISS-N4H5S2 | -0.05523 | 0.14813 | 0.71106 | 0.94815 |
| Thrombocytes | Blood Cell Counts | Gamma.MISS-N4H5S1F1 | 0.03178 | 0.14457 | 0.82702 | 0.94815 |
| Thrombocytes | Blood Cell Counts | Gamma.MISS-N4H5S1 | 0.09089 | 0.15045 | 0.54886 | 0.94815 |
| Thrombocytes | Blood Cell Counts | Beta.normal-N4H5S1 | -0.01650 | 0.16616 | 0.92143 | 0.94815 |
| Triglycerides | Biochemical Parameters and BMI | Gamma.MISS-N4H5S2 | 0.37857 | 0.07522 | 0.00000 | 0.00001 |
| Triglycerides | Biochemical Parameters and BMI | Gamma.MISS-N4H5S1 | -0.37129 | 0.07609 | 0.00000 | 0.00001 |
| Triglycerides | Biochemical Parameters and BMI | Beta.normal-N4H5S1 | -0.35565 | 0.08800 | 0.00010 | 0.00020 |
| Triglycerides | Biochemical Parameters and BMI | Beta.normal-N4H5S2 | 0.34710 | 0.08827 | 0.00015 | 0.00024 |
| Triglycerides | Biochemical Parameters and BMI | Gamma.MISS-N4H5S2F1 | 0.18745 | 0.07901 | 0.01888 | 0.02518 |
| Triglycerides | Biochemical Parameters and BMI | Gamma.MISS-N4H5S1F1 | 0.13492 | 0.08004 | 0.09387 | 0.10728 |
| Triglycerides | Biochemical Parameters and BMI | Gamma.MISS-N4H5 | -0.05399 | 0.07926 | 0.49673 | 0.49673 |
| Urea | Biochemical Parameters and BMI | Beta.normal-N4H5S2 | 0.07763 | 0.16432 | 0.63957 | 0.89665 |
| Urea | Biochemical Parameters and BMI | Gamma.MISS-N4H5S2 | 0.03117 | 0.15844 | 0.84498 | 0.89665 |
| Urea | Biochemical Parameters and BMI | Gamma.MISS-N4H5S1 | -0.02108 | 0.16133 | 0.89665 | 0.89665 |
| Urea | Biochemical Parameters and BMI | Beta.normal-N4H5S1 | -0.09637 | 0.16938 | 0.57302 | 0.89665 |
| Urea | Biochemical Parameters and BMI | Gamma.MISS-N4H5 | 0.06289 | 0.14895 | 0.67497 | 0.89665 |
| Urea | Biochemical Parameters and BMI | Gamma.MISS-N4H5S2F1 | 0.03929 | 0.15553 | 0.80175 | 0.89665 |
| Urea | Biochemical Parameters and BMI | Gamma.MISS-N4H5S1F1 | 0.03029 | 0.15432 | 0.84529 | 0.89665 |
